# On the front line of *Klebsiella pneumoniae* surface structures understanding: establishment of Fourier Transform Infrared (FT-IR) spectroscopy as a capsule typing method

**DOI:** 10.1101/554816

**Authors:** Carla Rodrigues, Clara Sousa, João Almeida Lopes, Ângela Novais, Luísa Peixe

## Abstract

Genomics-based population analysis of multidrug resistant (MDR) *Klebsiella pneumoniae* (*Kp*) motivated a renewed interest on capsule (K) types given their importance as evolutionary and virulence markers of clinically relevant strains. However, there is a gap between genotypic based predictions and information on capsular polysaccharide structure and composition. We used molecular genotypic, comparative genomics, biochemical and phenotypic data on the *cps* locus to support the usefulness of Fourier-Transform Infrared (FT-IR) spectroscopy as a phenotypic approach for K-type characterization and identification. The approach was validated with a collection of representative MDR *Kp* isolates from main lineages/Clonal Groups (CGs) involved in local or nationwide epidemics in 6 European and South American countries. FT-IR-based K-type assignments were compared with those obtained by genotypic methods and WGS (*cps* operon), and further complemented with data on the polysaccharide composition and structure of known K-types.

We demonstrate that our FT-IR-based spectroscopy approach can discriminate all 21 K-types identified with a resolution comparable (or even higher) to that provided by WGS, considered gold-standard methodology. Besides contributing to enlighten K-type diversity among a significant MDR *Kp* collection, the specific associations between certain K-types and *Kp* lineages identified in different geographic regions over time support the usefulness of our FT-IR-based approach for strain typing. Additionally, we demonstrate that FT-IR discriminatory ability is correlated with variation on the structure/composition of known K-types and, supported on WGS data, we were able to predict the sugar composition and chemical structure of new KL-types. Our data revealed an unprecedent resolution at a quick and low-cost rate of *Kp* K-types at the phenotypic level. Our FT-IR spectroscopy-based approach might be extremely useful not only as a cost-effective *Kp* typing tool, but also to improve our understanding on sugar-based coating structures of high relevance for strain evolution and host adaptation.

## INTRODUCTION

*Klebsiella pneumoniae* (*Kp*) is an encapsulated bacterium nowadays recognized as one of the most challenging human pathogens due to the increasing rates of mortality and morbidity associated with severe infections [hypervirulent (HV) strains] or with high rates of infections by strains resistant to multiple antibiotics [multidrug resistant (MDR) strains] ^1,2^. The capsule, an extracellular polysaccharide matrix, is one of the most striking virulence mechanisms considered essential for establishment of infection and for its protective effect against desiccation, phages and protists predation ^1^. Though it renders attractive properties as a target for vaccine development, the success of immunotherapeutic approaches depends on a complete understanding of bacterial surface structures circulating in the clinical setting ^3,4^. Whereas it is known that the expression of specific virulence factors and capsular (K) types (mainly K1 and K2) is related to severe infections caused by HV strains, much larger variation of capsular types has been described among clinical MDR strains, providing a higher resolution than multilocus sequence typing (MLST) and specific capsule-lineage associations that might be useful for typing ^5–7^. Variation on capsular (K antigen) and also other surface polysaccharides (such as O antigen) has been traditionally used for *Klebsiella* typing. In fact, serotyping, established as early as 1926 ^8^, allowed the recognition of 77 serologically distinct capsular types (K1-K82) and much less diverse O types (n=8, O1-O12) among the reference strains collection, deposited at Statens Serum Institute, Copenhagen, Denmark. ^9,10^. The chemical composition and structure of capsular types has been clarified essentially during the 1980’s for strains from the reference collection, but correlation with genomics data is recent and not always straightforward ^11^. The lack of practicability (complex and laborious) and availability (only at reference centres), and the insufficient coverage of serotyping led to its almost complete abandonment in the past decades ^1^.

Given the renewed interest in capsular polysaccharides to improve our understanding on the pathogenic potential of *Kp* and infection control strategies, several genotypic methods have recently been proposed to revive K-typing but there are several flaws that prevent their universal application and coverage. Molecular methods to infer K-type from genomics data such as RFLP of the *cps* locus (generating “C-patterns”) or PCR targeting specific K-types (e.g. K1, K2, K57) are technically demanding, have low coverage and/or are not suitable to detect variation in other sites of the locus ^12,13^. K-type prediction based on allelic variation of *loci* (e.g. *wzi* or *wzc*) within the capsule biosynthetic pathway (*cps* locus) constitute more rapid and simple approaches ^14,15^, but the characterization of the whole *cps* and *rfb* (O-antigen biosynthesis) clusters by whole genome sequencing (WGS) improved accessibility and precision, especially trough user-friendly web-based platforms (Kaptive) ^7,16^. These *in silico* studies and those that followed uncovered a series of novel surface locus (especially on *cps*), and at least 161 presumptive phenotypically distinct capsular types (designated KL to differentiate from the reference K-types) ^7,16–18^. More importantly, this data revealed the usefulness of K-variation as an epidemiological marker for strain subtyping, encouraging the development of reliable, fast and high-throughput tools for K-typing ^7,17,19–21^.

Fourier transform infrared (FT-IR) spectroscopy has been shown to detect surface phenotypic differences linked to a variable composition on glycan structures that form part of the O and K antigens, depending on the bacterial species ^22–25^. Considering that the capsule is the outermost structure of *Kp*, we hypothesize that FT-IR spectroscopy is a reliable tool to detect variations on *Kp* capsule composition, as observed for other capsulated bacteria ^26,27^. In this study, we combined genotypic, comparative genomics, biochemical and phenotypic data associated with the *cps* locus to assess the potential and robustness of FT-IR spectroscopy for identification of *Kp* capsular types. The high congruence established between capsular genotypic and phenotypic features of representative MDR *Kp* lineages circulating worldwide opens new avenues for a more comprehensive understanding on K-type variation and evolution and supports the potential of the methodology as an alternative fast and cost-effective *Kp* typing tool. The significance of the results for *K. pneumoniae* typing and for a better understanding of host-pathogen interactions is also discussed.

## RESULTS

### Molecular genotypic characterization of *Kp* K-antigen

There is a recent renewed interest on *Kp* capsular polysaccharides but the existing methodologies for K-tying are suboptimal and even with the development of WGS approaches, there is a lack of genotype – phenotype correlation. In this sense, we associated genotypic capsular data with FT-IR-based assignments and the biochemical composition of known capsule types to assess the suitability of FT-IR spectroscopy to identify known K-types or predict the composition of unknown K-types.

Our approach was validated on a collection of one hundred fifty-four well characterized MDR *Kp* isolates that had been involved in local or nationwide epidemics in different countries from Europe and South America spanning a long period of time (2002-2015). These isolates were selected to capture a diversity of capsule types encoded by main diverse *Kp* lineages from different clonal groups (CG) that had been involved in human clinical infections and in the expansion of extended-spectrum ß-lactamases (mainly CTX-M-15, SHV-12) and/or carbapenemases (mainly KPC-type, OXA-48, VIM) (**Table 1**). The performance of different cutting-edge genotypic K-typing methods will be compared with that provided by the phenotypic method FT-IR spectroscopy.

**Table 1.**
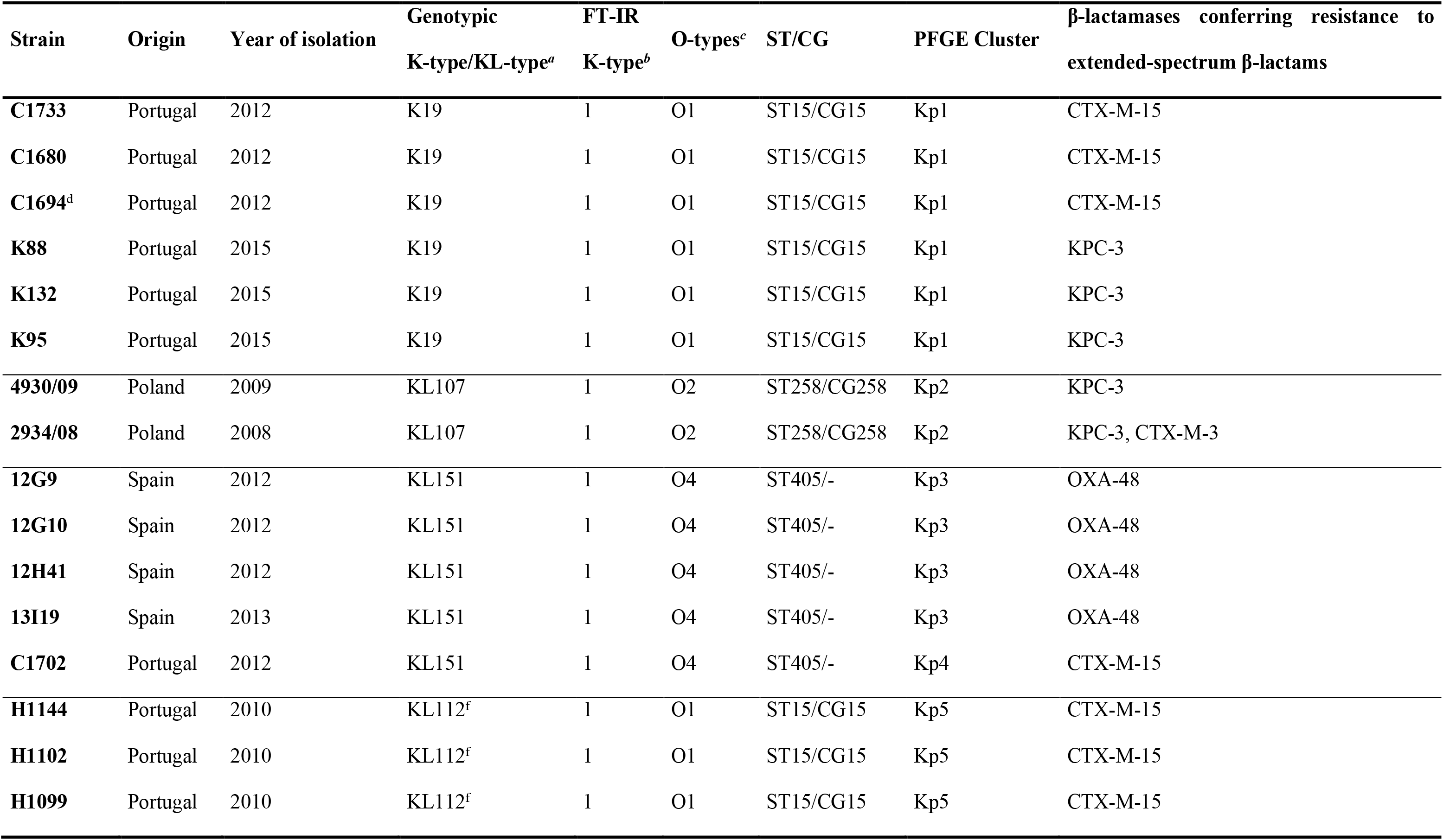
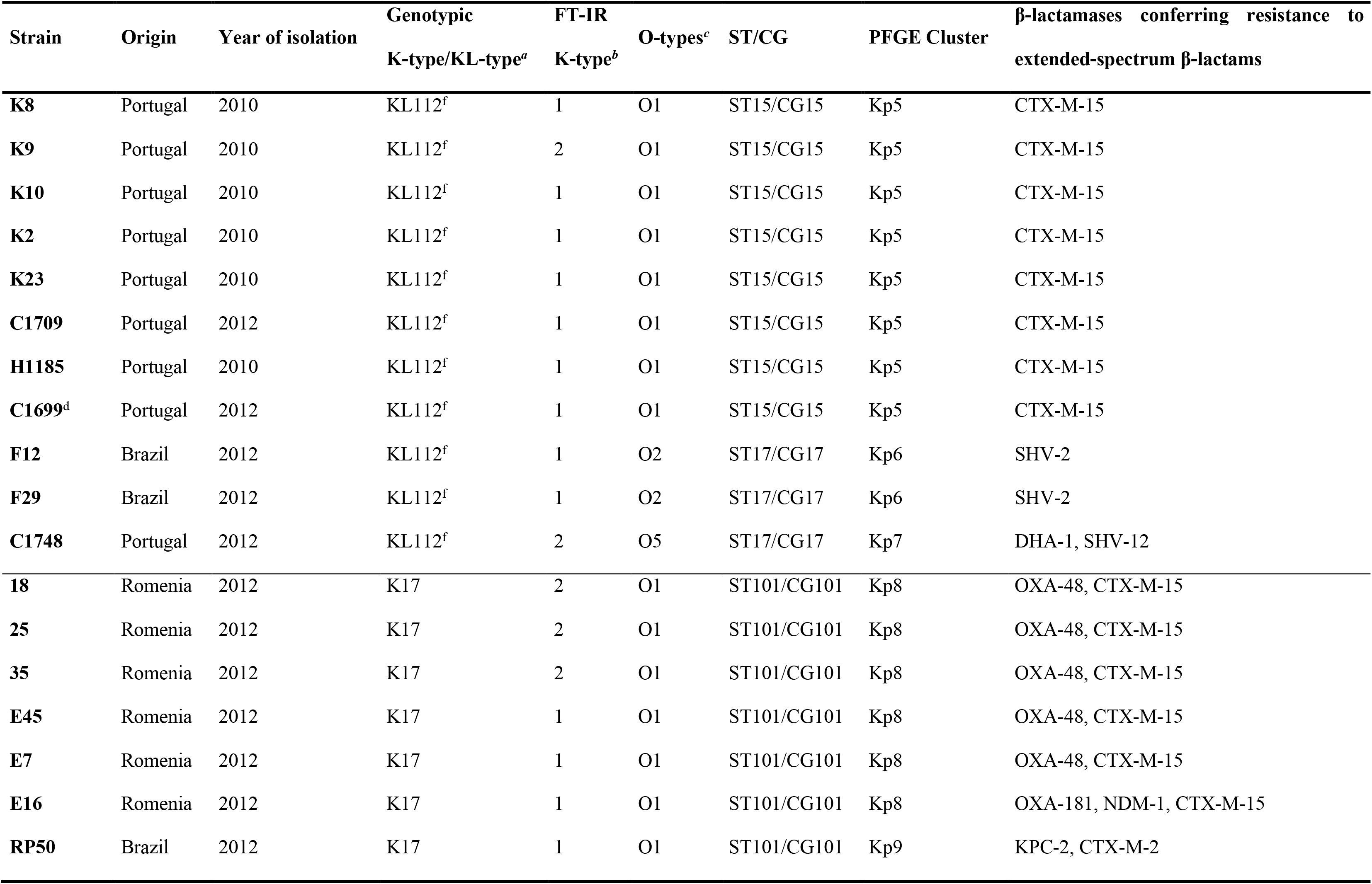
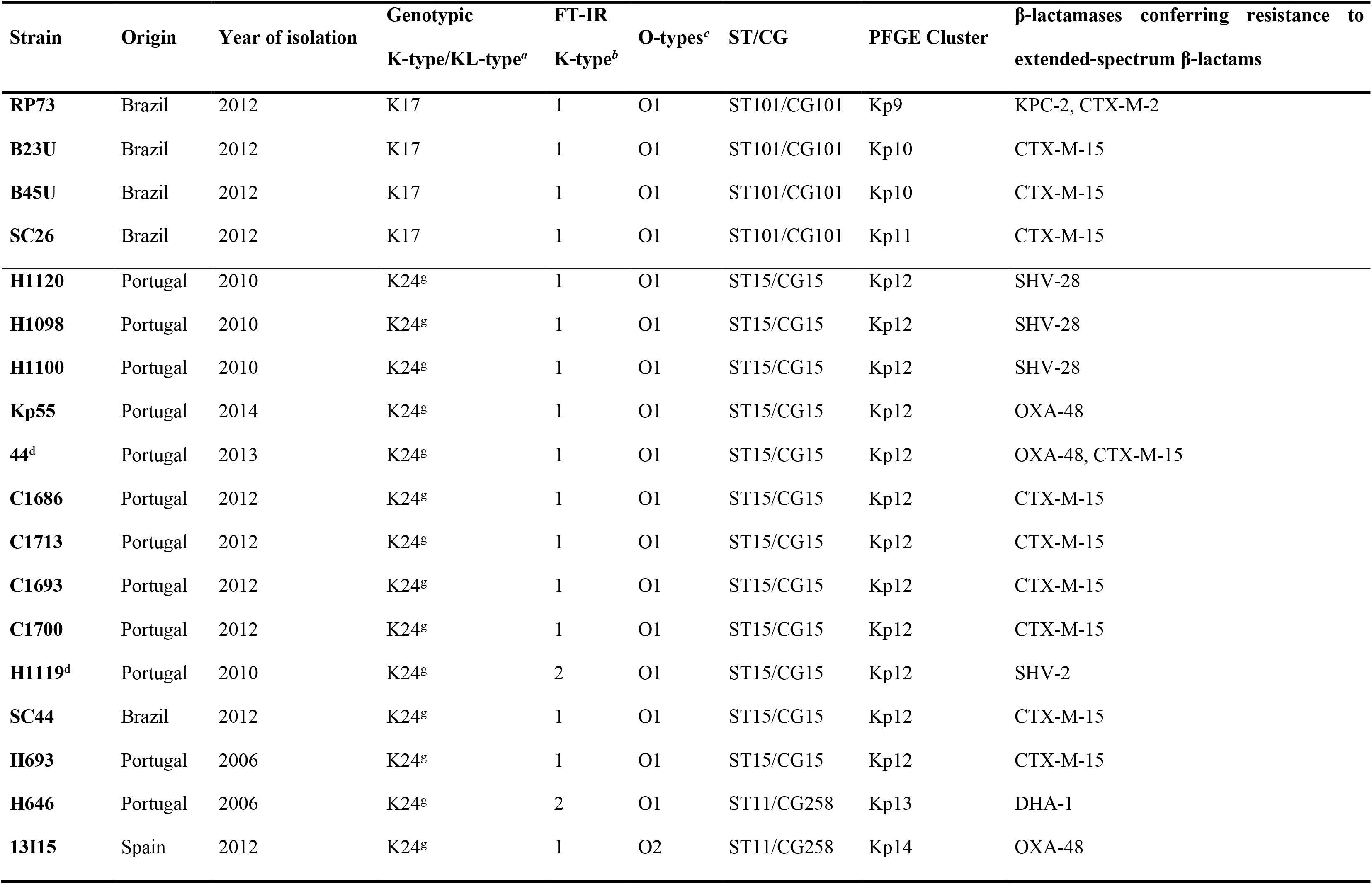
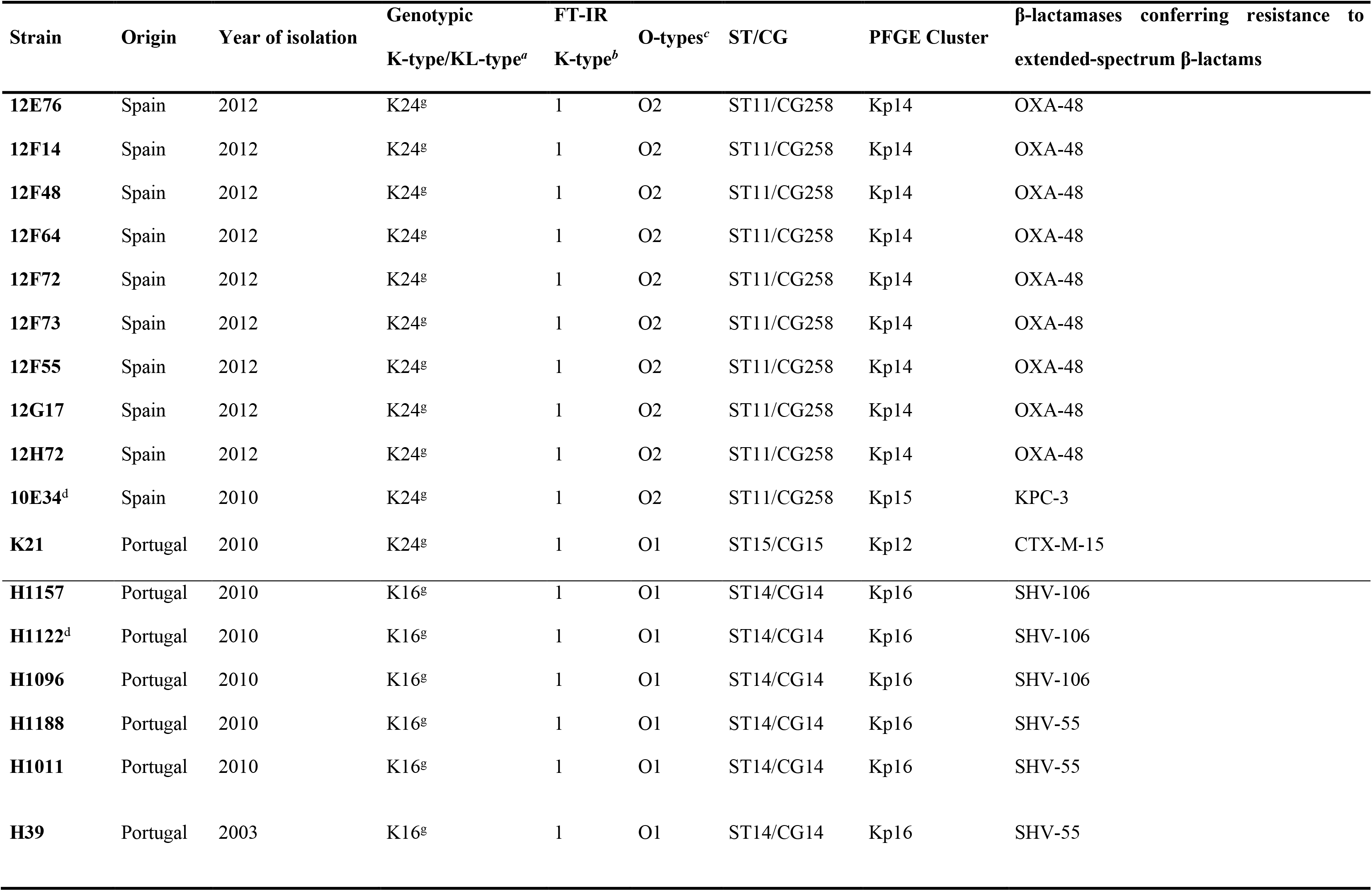
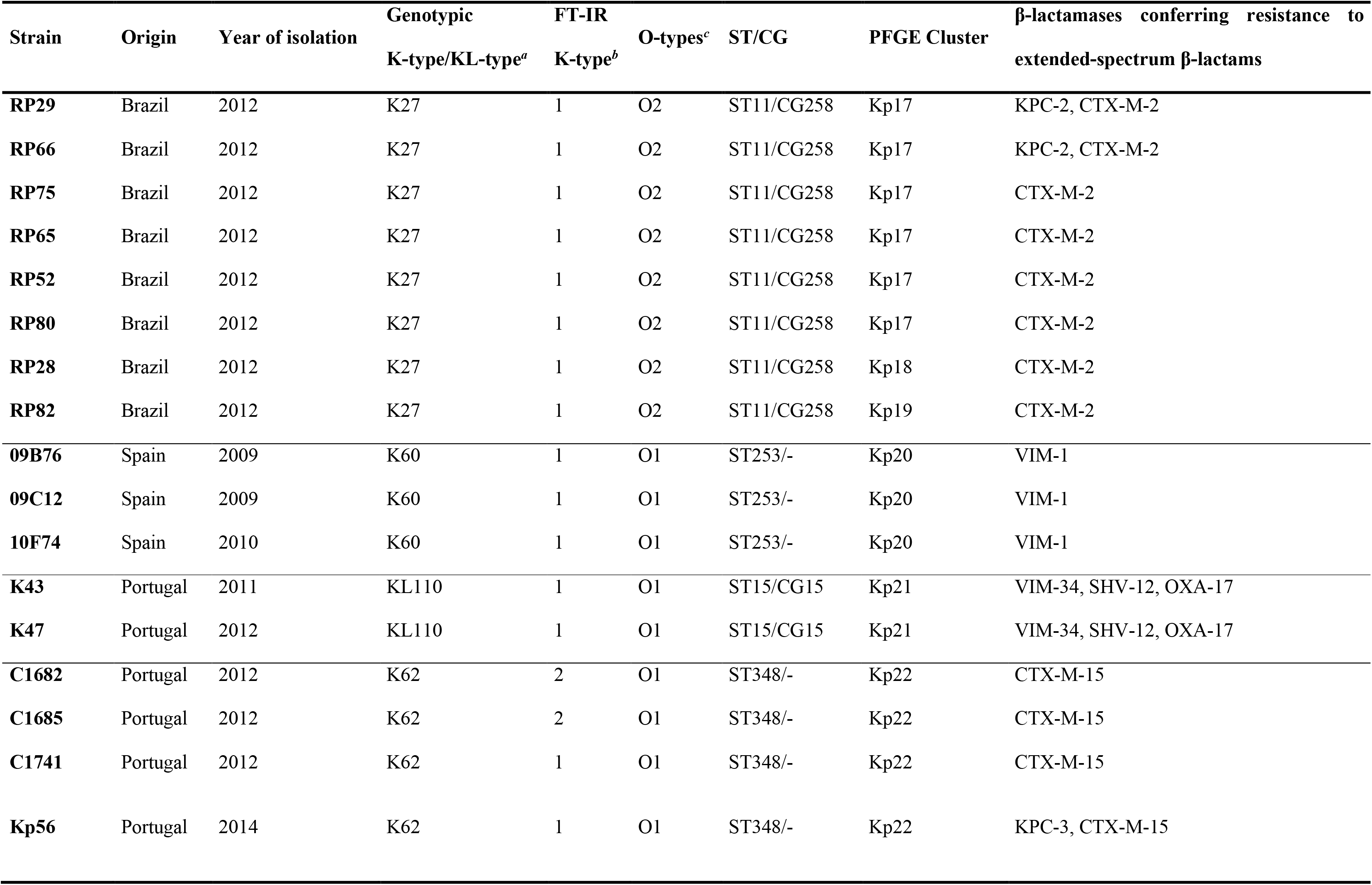
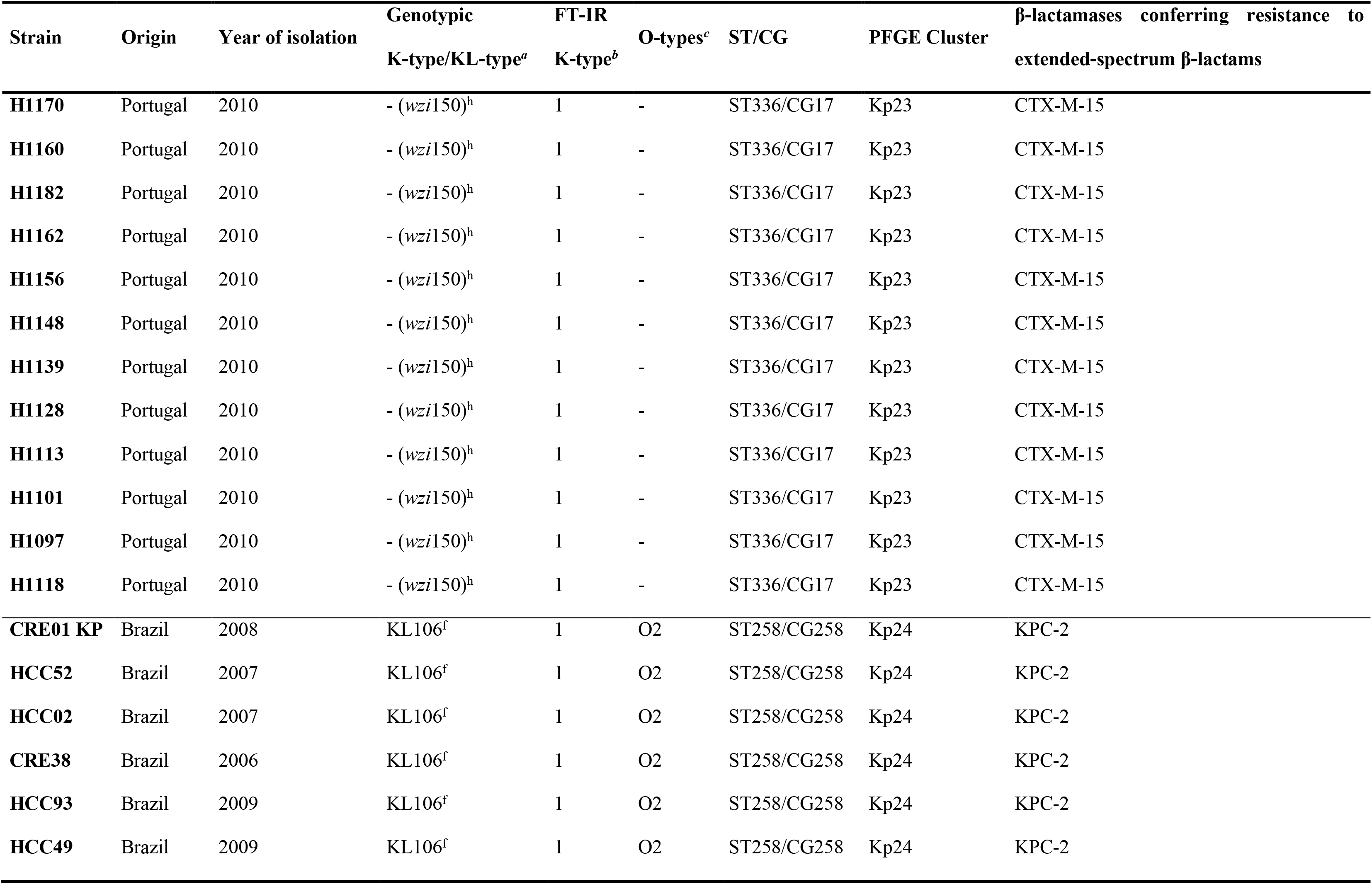
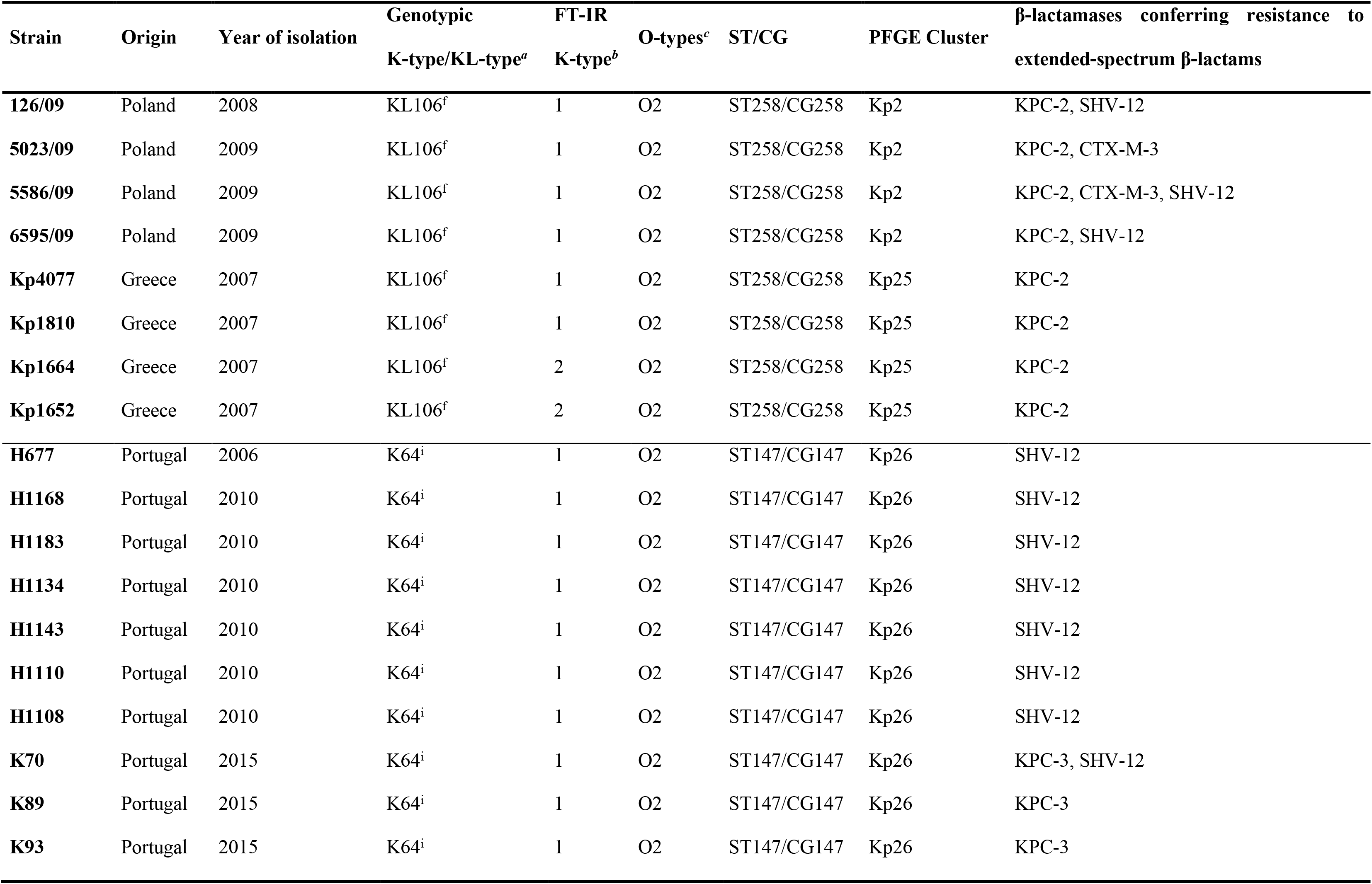
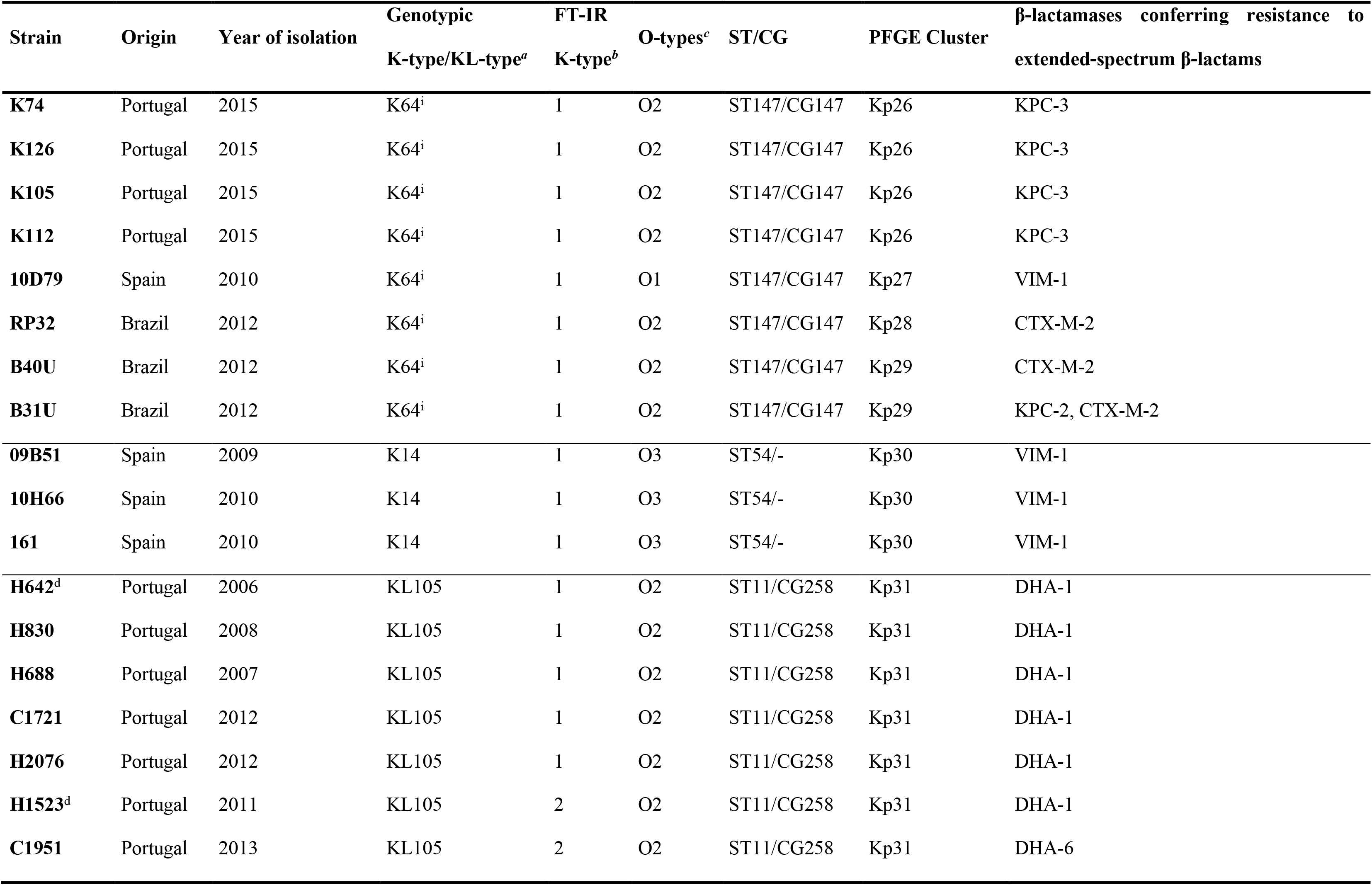
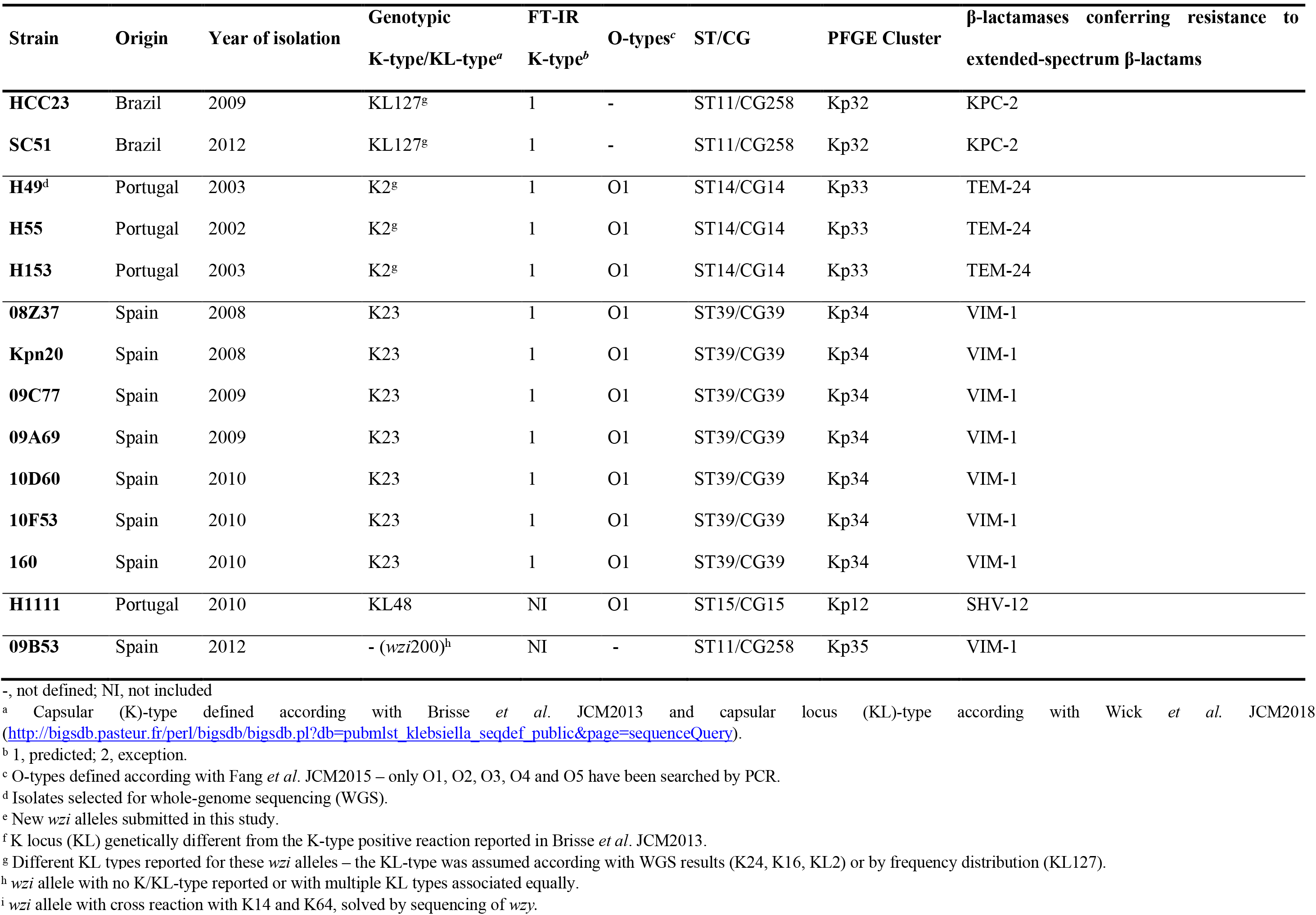
Epidemiological data and capsular characterization of international MDR *K. pneumoniae* clinical isolates analyzed in this study.

| Strain | Origin | Year of isolation | Genotypic<br>K-type/KL-type<sup>a</sup> | FT-IR<br>K-type<sup>b</sup> | O-types<sup>c</sup> | ST/CG | PFGE Cluster | β-lactamases conferring resistance to<br>extended-spectrum β-lactams |
| --- | --- | --- | --- | --- | --- | --- | --- | --- |
| **C1733** | Portugal | 2012 | K19 | 1 | O1 | ST15/CG15 | Kp1 | CTX-M-15 |
| **C1680** | Portugal | 2012 | K19 | 1 | O1 | ST15/CG15 | Kp1 | CTX-M-15 |
| **C1694<sup>d</sup>** | Portugal | 2012 | K19 | 1 | O1 | ST15/CG15 | Kp1 | CTX-M-15 |
| **K88** | Portugal | 2015 | K19 | 1 | O1 | ST15/CG15 | Kp1 | KPC-3 |
| **K132** | Portugal | 2015 | K19 | 1 | O1 | ST15/CG15 | Kp1 | KPC-3 |
| **K95** | Portugal | 2015 | K19 | 1 | O1 | ST15/CG15 | Kp1 | KPC-3 |
| **4930/09** | Poland | 2009 | KL107 | 1 | O2 | ST258/CG258 | Kp2 | KPC-3 |
| **2934/08** | Poland | 2008 | KL107 | 1 | O2 | ST258/CG258 | Kp2 | KPC-3, CTX-M-3 |
| **12G9** | Spain | 2012 | KL151 | 1 | O4 | ST405/- | Kp3 | OXA-48 |
| **12G10** | Spain | 2012 | KL151 | 1 | O4 | ST405/- | Kp3 | OXA-48 |
| **12H41** | Spain | 2012 | KL151 | 1 | O4 | ST405/- | Kp3 | OXA-48 |
| **13I19** | Spain | 2013 | KL151 | 1 | O4 | ST405/- | Kp3 | OXA-48 |
| **C1702** | Portugal | 2012 | KL151 | 1 | O4 | ST405/- | Kp4 | CTX-M-15 |
| **H1144** | Portugal | 2010 | KL112<sup>f</sup> | 1 | O1 | ST15/CG15 | Kp5 | CTX-M-15 |
| **H1102** | Portugal | 2010 | KL112<sup>f</sup> | 1 | O1 | ST15/CG15 | Kp5 | CTX-M-15 |
| **H1099** | Portugal | 2010 | KL112<sup>f</sup> | 1 | O1 | ST15/CG15 | Kp5 | CTX-M-15 |

| Strain | Origin | Year of isolation | Genotypic | FT-IR | O-types<sup>c</sup> | ST/CG | PFGE Cluster | β-lactamases conferring resistance to extended-spectrum β-lactams |
| --- | --- | --- | --- | --- | --- | --- | --- | --- |
|  |  |  | K-type/KL-type<sup>a</sup> | K-type<sup>b</sup> |  |  |  |  |
| **K8** | Portugal | 2010 | KL112<sup>f</sup> | 1 | O1 | ST15/CG15 | Kp5 | CTX-M-15 |
| **K9** | Portugal | 2010 | KL112<sup>f</sup> | 2 | O1 | ST15/CG15 | Kp5 | CTX-M-15 |
| **K10** | Portugal | 2010 | KL112<sup>f</sup> | 1 | O1 | ST15/CG15 | Kp5 | CTX-M-15 |
| **K2** | Portugal | 2010 | KL112<sup>f</sup> | 1 | O1 | ST15/CG15 | Kp5 | CTX-M-15 |
| **K23** | Portugal | 2010 | KL112<sup>f</sup> | 1 | O1 | ST15/CG15 | Kp5 | CTX-M-15 |
| **C1709** | Portugal | 2012 | KL112<sup>f</sup> | 1 | O1 | ST15/CG15 | Kp5 | CTX-M-15 |
| **H1185** | Portugal | 2010 | KL112<sup>f</sup> | 1 | O1 | ST15/CG15 | Kp5 | CTX-M-15 |
| **C1699<sup>d</sup>** | Portugal | 2012 | KL112<sup>f</sup> | 1 | O1 | ST15/CG15 | Kp5 | CTX-M-15 |
| **F12** | Brazil | 2012 | KL112<sup>f</sup> | 1 | O2 | ST17/CG17 | Kp6 | SHV-2 |
| **F29** | Brazil | 2012 | KL112<sup>f</sup> | 1 | O2 | ST17/CG17 | Kp6 | SHV-2 |
| **C1748** | Portugal | 2012 | KL112<sup>f</sup> | 2 | O5 | ST17/CG17 | Kp7 | DHA-1, SHV-12 |
| **18** | Romenia | 2012 | K17 | 2 | O1 | ST101/CG101 | Kp8 | OXA-48, CTX-M-15 |
| **25** | Romenia | 2012 | K17 | 2 | O1 | ST101/CG101 | Kp8 | OXA-48, CTX-M-15 |
| **35** | Romenia | 2012 | K17 | 2 | O1 | ST101/CG101 | Kp8 | OXA-48, CTX-M-15 |
| **E45** | Romenia | 2012 | K17 | 1 | O1 | ST101/CG101 | Kp8 | OXA-48, CTX-M-15 |
| **E7** | Romenia | 2012 | K17 | 1 | O1 | ST101/CG101 | Kp8 | OXA-48, CTX-M-15 |
| **E16** | Romenia | 2012 | K17 | 1 | O1 | ST101/CG101 | Kp8 | OXA-181, NDM-1, CTX-M-15 |
| **RP50** | Brazil | 2012 | K17 | 1 | O1 | ST101/CG101 | Kp9 | KPC-2, CTX-M-2 |

| Strain | Origin | Year of isolation | Genotypic<br>K-type/KL-type<sup>a</sup> | FT-IR<br>K-type<sup>b</sup> | O-types<sup>c</sup> | ST/CG | PFGE Cluster | β-lactamases conferring resistance to<br>extended-spectrum β-lactams |
| --- | --- | --- | --- | --- | --- | --- | --- | --- |
| **RP73** | Brazil | 2012 | K17 | 1 | O1 | ST101/CG101 | Kp9 | KPC-2, CTX-M-2 |
| **B23U** | Brazil | 2012 | K17 | 1 | O1 | ST101/CG101 | Kp10 | CTX-M-15 |
| **B45U** | Brazil | 2012 | K17 | 1 | O1 | ST101/CG101 | Kp10 | CTX-M-15 |
| **SC26** | Brazil | 2012 | K17 | 1 | O1 | ST101/CG101 | Kp11 | CTX-M-15 |
| **H1120** | Portugal | 2010 | K24<sup>g</sup> | 1 | O1 | ST15/CG15 | Kp12 | SHV-28 |
| **H1098** | Portugal | 2010 | K24<sup>g</sup> | 1 | O1 | ST15/CG15 | Kp12 | SHV-28 |
| **H1100** | Portugal | 2010 | K24<sup>g</sup> | 1 | O1 | ST15/CG15 | Kp12 | SHV-28 |
| **Kp55** | Portugal | 2014 | K24<sup>g</sup> | 1 | O1 | ST15/CG15 | Kp12 | OXA-48 |
| **44<sup>d</sup>** | Portugal | 2013 | K24<sup>g</sup> | 1 | O1 | ST15/CG15 | Kp12 | OXA-48, CTX-M-15 |
| **C1686** | Portugal | 2012 | K24<sup>g</sup> | 1 | O1 | ST15/CG15 | Kp12 | CTX-M-15 |
| **C1713** | Portugal | 2012 | K24<sup>g</sup> | 1 | O1 | ST15/CG15 | Kp12 | CTX-M-15 |
| **C1693** | Portugal | 2012 | K24<sup>g</sup> | 1 | O1 | ST15/CG15 | Kp12 | CTX-M-15 |
| **C1700** | Portugal | 2012 | K24<sup>g</sup> | 1 | O1 | ST15/CG15 | Kp12 | CTX-M-15 |
| **H1119<sup>d</sup>** | Portugal | 2010 | K24<sup>g</sup> | 2 | O1 | ST15/CG15 | Kp12 | SHV-2 |
| **SC44** | Brazil | 2012 | K24<sup>g</sup> | 1 | O1 | ST15/CG15 | Kp12 | CTX-M-15 |
| **H693** | Portugal | 2006 | K24<sup>g</sup> | 1 | O1 | ST15/CG15 | Kp12 | CTX-M-15 |
| **H646** | Portugal | 2006 | K24<sup>g</sup> | 2 | O1 | ST11/CG258 | Kp13 | DHA-1 |
| **13I15** | Spain | 2012 | K24<sup>g</sup> | 1 | O2 | ST11/CG258 | Kp14 | OXA-48 |

| Strain | Origin | Year of isolation | Genotypic | FT-IR | O-types<sup>c</sup> | ST/CG | PFGE Cluster | β-lactamases conferring resistance to extended-spectrum β-lactams |
| --- | --- | --- | --- | --- | --- | --- | --- | --- |
|  |  |  | K-type/KL-type<sup>a</sup> | K-type<sup>b</sup> |  |  |  |  |
| **12E76** | Spain | 2012 | K24<sup>g</sup> | 1 | O2 | ST11/CG258 | Kp14 | OXA-48 |
| **12F14** | Spain | 2012 | K24<sup>g</sup> | 1 | O2 | ST11/CG258 | Kp14 | OXA-48 |
| **12F48** | Spain | 2012 | K24<sup>g</sup> | 1 | O2 | ST11/CG258 | Kp14 | OXA-48 |
| **12F64** | Spain | 2012 | K24<sup>g</sup> | 1 | O2 | ST11/CG258 | Kp14 | OXA-48 |
| **12F72** | Spain | 2012 | K24<sup>g</sup> | 1 | O2 | ST11/CG258 | Kp14 | OXA-48 |
| **12F73** | Spain | 2012 | K24<sup>g</sup> | 1 | O2 | ST11/CG258 | Kp14 | OXA-48 |
| **12F55** | Spain | 2012 | K24<sup>g</sup> | 1 | O2 | ST11/CG258 | Kp14 | OXA-48 |
| **12G17** | Spain | 2012 | K24<sup>g</sup> | 1 | O2 | ST11/CG258 | Kp14 | OXA-48 |
| **12H72** | Spain | 2012 | K24<sup>g</sup> | 1 | O2 | ST11/CG258 | Kp14 | OXA-48 |
| **10E34<sup>d</sup>** | Spain | 2010 | K24<sup>g</sup> | 1 | O2 | ST11/CG258 | Kp15 | KPC-3 |
| **K21** | Portugal | 2010 | K24<sup>g</sup> | 1 | O1 | ST15/CG15 | Kp12 | CTX-M-15 |
| **H1157** | Portugal | 2010 | K16<sup>g</sup> | 1 | O1 | ST14/CG14 | Kp16 | SHV-106 |
| **H1122<sup>d</sup>** | Portugal | 2010 | K16<sup>g</sup> | 1 | O1 | ST14/CG14 | Kp16 | SHV-106 |
| **H1096** | Portugal | 2010 | K16<sup>g</sup> | 1 | O1 | ST14/CG14 | Kp16 | SHV-106 |
| **H1188** | Portugal | 2010 | K16<sup>g</sup> | 1 | O1 | ST14/CG14 | Kp16 | SHV-55 |
| **H1011** | Portugal | 2010 | K16<sup>g</sup> | 1 | O1 | ST14/CG14 | Kp16 | SHV-55 |
| **H39** | Portugal | 2003 | K16<sup>g</sup> | 1 | O1 | ST14/CG14 | Kp16 | SHV-55 |

| Strain | Origin | Year of isolation | Genotypic<br>K-type/KL-type<sup>a</sup> | FT-IR<br>K-type<sup>b</sup> | O-types<sup>c</sup> | ST/CG | PFGE Cluster | β-lactamases conferring resistance to<br>extended-spectrum β-lactams |
| --- | --- | --- | --- | --- | --- | --- | --- | --- |
| **RP29** | Brazil | 2012 | K27 | 1 | O2 | ST11/CG258 | Kp17 | KPC-2, CTX-M-2 |
| **RP66** | Brazil | 2012 | K27 | 1 | O2 | ST11/CG258 | Kp17 | KPC-2, CTX-M-2 |
| **RP75** | Brazil | 2012 | K27 | 1 | O2 | ST11/CG258 | Kp17 | CTX-M-2 |
| **RP65** | Brazil | 2012 | K27 | 1 | O2 | ST11/CG258 | Kp17 | CTX-M-2 |
| **RP52** | Brazil | 2012 | K27 | 1 | O2 | ST11/CG258 | Kp17 | CTX-M-2 |
| **RP80** | Brazil | 2012 | K27 | 1 | O2 | ST11/CG258 | Kp17 | CTX-M-2 |
| **RP28** | Brazil | 2012 | K27 | 1 | O2 | ST11/CG258 | Kp18 | CTX-M-2 |
| **RP82** | Brazil | 2012 | K27 | 1 | O2 | ST11/CG258 | Kp19 | CTX-M-2 |
| **09B76** | Spain | 2009 | K60 | 1 | O1 | ST253/- | Kp20 | VIM-1 |
| **09C12** | Spain | 2009 | K60 | 1 | O1 | ST253/- | Kp20 | VIM-1 |
| **10F74** | Spain | 2010 | K60 | 1 | O1 | ST253/- | Kp20 | VIM-1 |
| **K43** | Portugal | 2011 | KL110 | 1 | O1 | ST15/CG15 | Kp21 | VIM-34, SHV-12, OXA-17 |
| **K47** | Portugal | 2012 | KL110 | 1 | O1 | ST15/CG15 | Kp21 | VIM-34, SHV-12, OXA-17 |
| **C1682** | Portugal | 2012 | K62 | 2 | O1 | ST348/- | Kp22 | CTX-M-15 |
| **C1685** | Portugal | 2012 | K62 | 2 | O1 | ST348/- | Kp22 | CTX-M-15 |
| **C1741** | Portugal | 2012 | K62 | 1 | O1 | ST348/- | Kp22 | CTX-M-15 |
| **Kp56** | Portugal | 2014 | K62 | 1 | O1 | ST348/- | Kp22 | KPC-3, CTX-M-15 |

| Strain | Origin | Year of isolation | Genotypic | FT-IR | O-types<sup>c</sup> | ST/CG | PFGE Cluster | β-lactamases conferring resistance to extended-spectrum β-lactams |
| --- | --- | --- | --- | --- | --- | --- | --- | --- |
|  |  |  | K-type/KL-type<sup>a</sup> | K-type<sup>b</sup> |  |  |  |  |
| **H1170** | Portugal | 2010 | - (*wzi150*)<sup>h</sup> | 1 | - | ST336/CG17 | Kp23 | CTX-M-15 |
| **H1160** | Portugal | 2010 | - (*wzi150*)<sup>h</sup> | 1 | - | ST336/CG17 | Kp23 | CTX-M-15 |
| **H1182** | Portugal | 2010 | - (*wzi150*)<sup>h</sup> | 1 | - | ST336/CG17 | Kp23 | CTX-M-15 |
| **H1162** | Portugal | 2010 | - (*wzi150*)<sup>h</sup> | 1 | - | ST336/CG17 | Kp23 | CTX-M-15 |
| **H1156** | Portugal | 2010 | - (*wzi150*)<sup>h</sup> | 1 | - | ST336/CG17 | Kp23 | CTX-M-15 |
| **H1148** | Portugal | 2010 | - (*wzi150*)<sup>h</sup> | 1 | - | ST336/CG17 | Kp23 | CTX-M-15 |
| **H1139** | Portugal | 2010 | - (*wzi150*)<sup>h</sup> | 1 | - | ST336/CG17 | Kp23 | CTX-M-15 |
| **H1128** | Portugal | 2010 | - (*wzi150*)<sup>h</sup> | 1 | - | ST336/CG17 | Kp23 | CTX-M-15 |
| **H1113** | Portugal | 2010 | - (*wzi150*)<sup>h</sup> | 1 | - | ST336/CG17 | Kp23 | CTX-M-15 |
| **H1101** | Portugal | 2010 | - (*wzi150*)<sup>h</sup> | 1 | - | ST336/CG17 | Kp23 | CTX-M-15 |
| **H1097** | Portugal | 2010 | - (*wzi150*)<sup>h</sup> | 1 | - | ST336/CG17 | Kp23 | CTX-M-15 |
| **H1118** | Portugal | 2010 | - (*wzi150*)<sup>h</sup> | 1 | - | ST336/CG17 | Kp23 | CTX-M-15 |
| **CRE01 KP** | Brazil | 2008 | KL106<sup>f</sup> | 1 | O2 | ST258/CG258 | Kp24 | KPC-2 |
| **HCC52** | Brazil | 2007 | KL106<sup>f</sup> | 1 | O2 | ST258/CG258 | Kp24 | KPC-2 |
| **HCC02** | Brazil | 2007 | KL106<sup>f</sup> | 1 | O2 | ST258/CG258 | Kp24 | KPC-2 |
| **CRE38** | Brazil | 2006 | KL106<sup>f</sup> | 1 | O2 | ST258/CG258 | Kp24 | KPC-2 |
| **HCC93** | Brazil | 2009 | KL106<sup>f</sup> | 1 | O2 | ST258/CG258 | Kp24 | KPC-2 |
| **HCC49** | Brazil | 2009 | KL106<sup>f</sup> | 1 | O2 | ST258/CG258 | Kp24 | KPC-2 |
| **126/09** | Poland | 2008 | KL106<sup>f</sup> | 1 | O2 | ST258/CG258 | Kp2 | KPC-2, SHV-12 |
| **5023/09** | Poland | 2009 | KL106<sup>f</sup> | 1 | O2 | ST258/CG258 | Kp2 | KPC-2, CTX-M-3 |
| **5586/09** | Poland | 2009 | KL106<sup>f</sup> | 1 | O2 | ST258/CG258 | Kp2 | KPC-2, CTX-M-3, SHV-12 |
| **6595/09** | Poland | 2009 | KL106<sup>f</sup> | 1 | O2 | ST258/CG258 | Kp2 | KPC-2, SHV-12 |
| **Kp4077** | Greece | 2007 | KL106<sup>f</sup> | 1 | O2 | ST258/CG258 | Kp25 | KPC-2 |
| **Kp1810** | Greece | 2007 | KL106<sup>f</sup> | 1 | O2 | ST258/CG258 | Kp25 | KPC-2 |
| **Kp1664** | Greece | 2007 | KL106<sup>f</sup> | 2 | O2 | ST258/CG258 | Kp25 | KPC-2 |
| **Kp1652** | Greece | 2007 | KL106<sup>f</sup> | 2 | O2 | ST258/CG258 | Kp25 | KPC-2 |
| **H677** | Portugal | 2006 | K64<sup>i</sup> | 1 | O2 | ST147/CG147 | Kp26 | SHV-12 |
| **H1168** | Portugal | 2010 | K64<sup>i</sup> | 1 | O2 | ST147/CG147 | Kp26 | SHV-12 |
| **H1183** | Portugal | 2010 | K64<sup>i</sup> | 1 | O2 | ST147/CG147 | Kp26 | SHV-12 |
| **H1134** | Portugal | 2010 | K64<sup>i</sup> | 1 | O2 | ST147/CG147 | Kp26 | SHV-12 |
| **H1143** | Portugal | 2010 | K64<sup>i</sup> | 1 | O2 | ST147/CG147 | Kp26 | SHV-12 |
| **H1110** | Portugal | 2010 | K64<sup>i</sup> | 1 | O2 | ST147/CG147 | Kp26 | SHV-12 |
| **H1108** | Portugal | 2010 | K64<sup>i</sup> | 1 | O2 | ST147/CG147 | Kp26 | SHV-12 |
| **K70** | Portugal | 2015 | K64<sup>i</sup> | 1 | O2 | ST147/CG147 | Kp26 | KPC-3, SHV-12 |
| **K89** | Portugal | 2015 | K64<sup>i</sup> | 1 | O2 | ST147/CG147 | Kp26 | KPC-3 |
| **K93** | Portugal | 2015 | K64<sup>i</sup> | 1 | O2 | ST147/CG147 | Kp26 | KPC-3 |

| Strain | Origin | Year of isolation | Genotypic | FT-IR | O-types<sup>c</sup> | ST/CG | PFGE Cluster | β-lactamases conferring resistance to extended-spectrum β-lactams |
| --- | --- | --- | --- | --- | --- | --- | --- | --- |
|  |  |  | K-type/KL-type<sup>a</sup> | K-type<sup>b</sup> |  |  |  |  |
| **K74** | Portugal | 2015 | K64<sup>i</sup> | 1 | O2 | ST147/CG147 | Kp26 | KPC-3 |
| **K126** | Portugal | 2015 | K64<sup>i</sup> | 1 | O2 | ST147/CG147 | Kp26 | KPC-3 |
| **K105** | Portugal | 2015 | K64<sup>i</sup> | 1 | O2 | ST147/CG147 | Kp26 | KPC-3 |
| **K112** | Portugal | 2015 | K64<sup>i</sup> | 1 | O2 | ST147/CG147 | Kp26 | KPC-3 |
| **10D79** | Spain | 2010 | K64<sup>i</sup> | 1 | O1 | ST147/CG147 | Kp27 | VIM-1 |
| **RP32** | Brazil | 2012 | K64<sup>i</sup> | 1 | O2 | ST147/CG147 | Kp28 | CTX-M-2 |
| **B40U** | Brazil | 2012 | K64<sup>i</sup> | 1 | O2 | ST147/CG147 | Kp29 | CTX-M-2 |
| **B31U** | Brazil | 2012 | K64<sup>i</sup> | 1 | O2 | ST147/CG147 | Kp29 | KPC-2, CTX-M-2 |
| **09B51** | Spain | 2009 | K14 | 1 | O3 | ST54/- | Kp30 | VIM-1 |
| **10H66** | Spain | 2010 | K14 | 1 | O3 | ST54/- | Kp30 | VIM-1 |
| **161** | Spain | 2010 | K14 | 1 | O3 | ST54/- | Kp30 | VIM-1 |
| **H642<sup>d</sup>** | Portugal | 2006 | KL105 | 1 | O2 | ST11/CG258 | Kp31 | DHA-1 |
| **H830** | Portugal | 2008 | KL105 | 1 | O2 | ST11/CG258 | Kp31 | DHA-1 |
| **H688** | Portugal | 2007 | KL105 | 1 | O2 | ST11/CG258 | Kp31 | DHA-1 |
| **C1721** | Portugal | 2012 | KL105 | 1 | O2 | ST11/CG258 | Kp31 | DHA-1 |
| **H2076** | Portugal | 2012 | KL105 | 1 | O2 | ST11/CG258 | Kp31 | DHA-1 |
| **H1523<sup>d</sup>** | Portugal | 2011 | KL105 | 2 | O2 | ST11/CG258 | Kp31 | DHA-1 |
| **C1951** | Portugal | 2013 | KL105 | 2 | O2 | ST11/CG258 | Kp31 | DHA-6 |
| HCC23 | Brazil | 2009 | KL127<sup>g</sup> | 1 | - | ST11/CG258 | Kp32 | KPC-2 |
| SC51 | Brazil | 2012 | KL127<sup>g</sup> | 1 | - | ST11/CG258 | Kp32 | KPC-2 |
| H49<sup>d</sup> | Portugal | 2003 | K2<sup>g</sup> | 1 | O1 | ST14/CG14 | Kp33 | TEM-24 |
| H55 | Portugal | 2002 | K2<sup>g</sup> | 1 | O1 | ST14/CG14 | Kp33 | TEM-24 |
| H153 | Portugal | 2003 | K2<sup>g</sup> | 1 | O1 | ST14/CG14 | Kp33 | TEM-24 |
| 08Z37 | Spain | 2008 | K23 | 1 | O1 | ST39/CG39 | Kp34 | VIM-1 |
| Kpn20 | Spain | 2008 | K23 | 1 | O1 | ST39/CG39 | Kp34 | VIM-1 |
| 09C77 | Spain | 2009 | K23 | 1 | O1 | ST39/CG39 | Kp34 | VIM-1 |
| 09A69 | Spain | 2009 | K23 | 1 | O1 | ST39/CG39 | Kp34 | VIM-1 |
| 10D60 | Spain | 2010 | K23 | 1 | O1 | ST39/CG39 | Kp34 | VIM-1 |
| 10F53 | Spain | 2010 | K23 | 1 | O1 | ST39/CG39 | Kp34 | VIM-1 |
| 160 | Spain | 2010 | K23 | 1 | O1 | ST39/CG39 | Kp34 | VIM-1 |
| H1111 | Portugal | 2010 | KL48 | NI | O1 | ST15/CG15 | Kp12 | SHV-12 |
| 09B53 | Spain | 2012 | -(wzi200)<sup>h</sup> | NI | - | ST11/CG258 | Kp35 | VIM-1 |

-, not defined; NI, not included
<sup>a</sup> Capsular (K)-type defined according with Brisse *et al.* JCM2013 and capsular locus (KL)-type according with Wick *et al.* JCM2018 (http://bigsd.b.pasteur.fr/perl/bigsd/bigsd.pl?db=pubmlst_klebsiella_seqdef_public&page=sequenceQuery).
<sup>b</sup> 1, predicted; 2, exception.
<sup>c</sup> O-types defined according with Fang *et al.* JCM2015 – only O1, O2, O3, O4 and O5 have been searched by PCR.
<sup>d</sup> Isolates selected for whole-genome sequencing (WGS).
<sup>e</sup> New *wzi* alleles submitted in this study.
<sup>f</sup> K locus (KL) genetically different from the K-type positive reaction reported in Brisse *et al.* JCM2013.
<sup>g</sup> Different KL types reported for these *wzi* alleles – the KL-type was assumed according with WGS results (K24, K16, KL2) or by frequency distribution (KL127).
<sup>h</sup> *wzi* allele with no K/KL-type reported or with multiple KL types associated equally. <sup>i</sup> *wzi* allele with cross reaction with K14 and K64, solved by sequencing of *wzy*.

#### Capsular assignment based on the genotypic marker

wzi. First, capsular types were inferred by sequence comparison of a discriminatory molecular marker (*wzi*) ^14^. Considering this molecular genotypic approach, the collection analysed encompassed 22 different *wzi* alleles, four of which (*wzi*89, *wzi*200-202) were newly described and deposited at the BIGSdb-*Kp* Pasteur database (http://bigsdb.pasteur.fr/klebsiella/klebsiella.html), and described meanwhile in other studies ^16^. According to the BIGSdb-*Kp* database, these *wzi* alleles are presumptively associated with certain K-types (positive reaction with the sera from reference K-types) and/or KL-types (which stands for the *cps* locus types obtained by WGS data, when available) (**Table 2**). In thirteen of them, the *wzi* allele was unequivocally associated with one unique K-type and/or KL-type. However, prediction of K-type was not always straightforward since: i) 6 *wzi* alleles were linked to more than one K-type/KL-type; ii) 2 *wzi* alleles (*wzi*29, *wzi*93) were linked to discordant K-type/KL-type; and iii) 1 *wzi* allele (*wzi*200) has no K/KL-type attributed (Table 2).

**Table 2.** K/KL-types predicted based on genotypic, genomic and phenotypic methods.

| Prediction based on *wzi* |  |  | Prediction based on WGS/Kaptive<sup>c</sup> | Prediction based on FT-IR |
| --- | --- | --- | --- | --- |
| *wzi* allele | associated K-type<sup>a</sup> | associated KL type<sup>b</sup> |  |  |
| 2 | K2 | KL2.KL30 | **KL2** | K2 |
| 14 | K14 | KL14 | KL14 | K14 |
| 16 | K16 | KL16.KL143 | **KL16** | K16 |
| 19 | K19 | KL19 | **KL19** | K19 |
| 24 | K24 | KL24.KL54.KL55 | **KL24** | K24<sup>d</sup> |
| 27 | K27 | KL27 | KL27 | K27 |
| 29 | K41 | KL106 | KL106 | KL106<sup>d</sup> |
| 64 | K14.K64 | KL64 | KL64 | KL64 |
| 75 | - | KL105 | **KL105** | KL105<sup>d</sup> |
| 83 | K23 | KL23 | KL23 | K23 |
| 89 | - | KL110 | KL110 | KL110 |
| 93 | K60 | KL112 | **KL112** | KL112<sup>d</sup> |
| 94 | - | KL62 | KL62 | K62<sup>d</sup> |
| 101 | K24 | KL24 | KL24 | K24 |
| 137 | K17 | KL17 | KL17 | K17<sup>d</sup> |
| 143 | - | KL151 | KL151 | KL151 |
| 150 | - | KL163.KL27.KL46 | - | -<sup>e</sup> |
| 151 | - | KL48 | KL48 | KL48 |
| 154 | - | KL107 | KL107 | KL107 |
| 200 | - | - | - | - |
| 201 | - | KL60 | KL60 | KL60 |
| 202 | - | KL127.KL155 | KL127 | KL127 |

-, not defined
<sup>a</sup> known associated K-types (serological reaction tested by Brisse *et al.* 2013)
<sup>b</sup> associated *cps* cluster type (KL stands for K locus)
<sup>c</sup> in bold the KL-type was confirmed by WGS results, the remaining are based in frequency distribution (KL127).
<sup>d</sup> presence of outliers outside the main cluster.
<sup>e</sup> according to FT-IR results we can discard the possibility of being a KL27.

#### *Capsular prediction based on* wzy *or epidemiological data*

Some of these uncertain K-types were additionally defined by sequencing of another molecular marker (*wzy*), by analysis of available epidemiological data (where the most frequently reported K-type for a given ST was considered), or by WGS and Kaptive (see below) (**Table 2**) ^7,15,16^. Sequencing of *wzy* allowed distinguishing K14/K64 predicted by *wzi*64, whereas the epidemiological information supported the prediction of capsule types K2, K16, K24, KL106, KL112 and KL127. With this approach, a total of 19 different K-types were predicted by *wzi/wzy* sequencing, that varied in frequency between 0.7% to 17.7% (**Table 1**). Twelve of them belong to K-types serologically defined (K2, K14, K16, K17, K19, K23, K24, K27, KL48, KL60, KL62 and K64) and 7 are K-types presumptively associated with a new composition/structure (KL105, KL106, KL107, KL110, KL112, KL127 and KL151) (**Table 1**). However, interpretation of K/KL-types is not straightforward and/or needs to be supported by epidemiological information.

It is of interest to highlight that most K-types identified in this collection were specifically and uniquely associated with evolutionarily related strains from different countries and recovered from large periods of time (**Table 1**). Some of them correspond to well-established clades from CG11 ^7,21,28^, CG15 or CG14 ^7,20,29^ and CG258 ^19,30^ identified in previous studies. Occasionally, the same K-type was observed in different clones (e.g. K24 in ST11 and ST15, or K64 in ST11 and ST147) (**Table 1**).

### Molecular genotypic characterization of *Kp* O-antigen

Considering that the O antigen can in some isolates protrude to the bacterial cell surface depending on the amount and type of the capsule, we cannot despise its potential contribution to the biochemical make-up of the bacterial cell surface. In this sense, a molecular genotypic PCR-based approach was used to identify the most frequent O-types previously recognized among *Kp* clinical isolates (O1, O2, O3, O4 and O5) ^10^. As expected, much less diversity of O-types characterizes the collection analyzed. Most isolates belong to O1 (44.8%, 69/154) and O2 (39.6%, 61/154), followed by O4 (3.2%, 5/154), O3 (1.9%, 3/154) and O5 (0.7%, 1/154), while 9.7% (15/154) of the isolates were not typed with the primers used. We also observed that isolates belonging to the same clone and exhibiting a given capsular type had the same O-type, with very few exceptions. It is of note that evolutionarily related isolates belonging to the same or closely related clonal group such as ST15 and ST14 from CG15/CG14 or ST11 and ST258 from CG258, exhibited the same O-type (O1 for CG15/CG14 and O2 for CG258) (**Table 1**).

### Comparative genomics of *cps* gene clusters

Full discrimination of capsular types has been possible by sequencing of the whole *cps* cluster by WGS, that supported the assignment of KL-types for which the composition/structure by biochemical procedures is still unreported, some of which being especially associated with worldwide disseminated clones such as ST258 or ST15. Furthermore, it is well known that changes in sites of the *cps* locus other than *wzi* or *wzy* can influence the final capsule composition. To provide full K-typing resolution and support FT-IR-based phenotypic assignments, we performed a comparative genomics analysis of *cps* clusters of the 19 *wzi*-defined K-/KL-types. We used *cps* available at the NCBI GenBank database for strains of the reference *Kp* collection and also performed *de novo* genome sequencing of 9 isolates from this study (**Figure 1**). The isolates harbouring *wzi*150 and *wzi*200 were excluded since the K/KL-type was unclear or unknown (**Table 2**).

**Figure 1.**
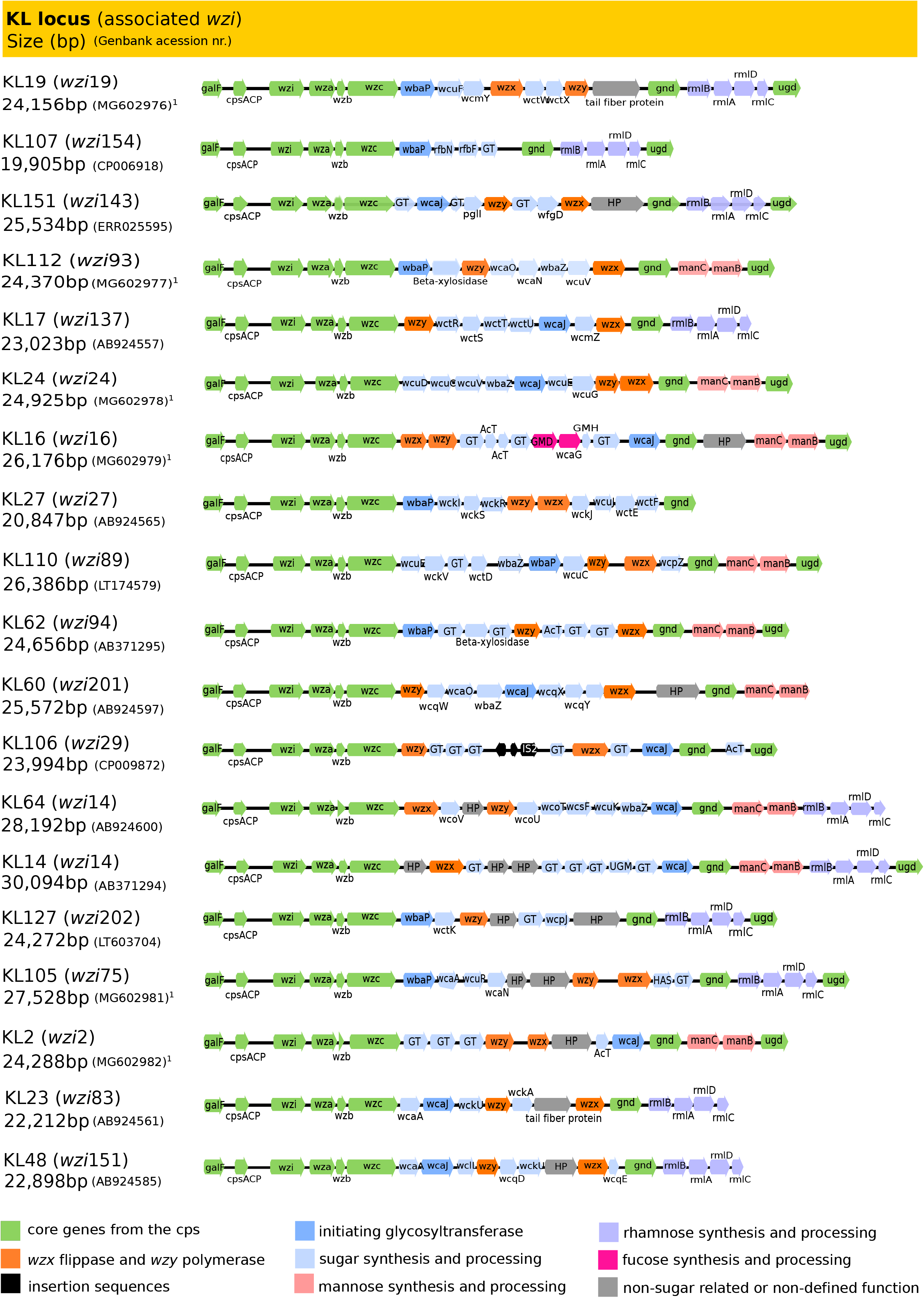
Representation of the *cps* genetic cluster identified in this study. Legend: ^1^, this study; HP, hypothetical protein; GT, glycosyltransferase; AcT, acyltransferase, GMD, GDP-mannose-4,6-dehydratase; GMH, GDP-mannose mannosyl hydrolase; GH, glycolsylhydrolase; HAS, hyaluronan synthase; UGM, UDP-galactopyranose mutase.

#### WGS-based K-type assignments

The *cps* clusters presented a variable size (20 – 30 Kb) and were delimited by the conserved *galF* (encodes an UTP–glucose-1-phosphate uridylyltransferase responsible for the synthesis of UDP-G-glucose) and, in most cases, *ugd* (encodes a UDP-glucose 6-dehydrogenase, responsible for the formation of UDP-D-glucoronate) genes, and possessed a series of other highly conserved genes at the 5’ of the locus. Coding sequences (between 16 and 25) were represented and compared using Geneious R10 software (Biomatters Ltd, Auckland, New Zealand) (Figure 1). Each of the *cps* locus represented contains a unique combination of genes that is predictive of 19 different K-/KL-types. Whole *cps*-based typing allowed: i) confirmation of K-type predictions based on *wzi* sequencing and available epidemiological data, when there was insufficient precision (e.g. K2, K16 or K24); ii) clarifying discrepancies between K-type and KL-types (e.g. K-type KL112), and iii) unveiling small genetic differences in other sites of the locus, that were subsequently correlated with phenotypic changes detected by FT-IR (KL105 isolates, Figure 4, see below). Thus, our data confirms that WGS provides a higher resolution for *Kp* K-typing, but also that genomics data might be insufficient to precisely predict final capsule composition. It is also of remark that *cps* sequenced in this study were identical (100-99%) to those reported in isolates from the reference collection or previously deposited in public databases (Table S1) (data not shown).

#### *Analysis of* cps *genes involved in sugar synthesis*

In a close analysis of all *cps* clusters, special attention was paid to the presence of genes associated with the synthesis of particular sugars: i) initial glycosyltransferases responsible for triggering capsule synthesis. The *wbaP* (encoding an undecaprenyl phosphate galactose transferase) and *wcaJ* (encoding an undecaprenyl-phosphate glucose-1-phosphate) genes were detected 8 or 9 of the *cps* clusters, respectively. The corresponding proteins revealed a high degree of homology (∼70% identity) (data not shown) and are, respectively, predictive of the presence of galactose or glucose on the repeat unit ^11^; ii) genes responsible for the synthesis of L-fucose (*gmd* and *wcaG*; n=1/19, 5%), GDP-D-mannose (*manCB*; n=9/19, 47%) and UDP-L-rhamnose (*rmlBADC*; n=10/19, 53%) were identified in the variable regions between *wzc* and *gnd* or between *gnd* and *ugd*. A series of other genes encoding putative non-initial glycosyltransferases, modifying enzymes (acetyltransferases, pyruvyl transferases, glycosyl hydrolases), insertion sequences (IS) or hypothetical proteins were also detected (**Figure 1**). These genotypic data supported correlations established with the presence of different sugars in the final capsule polysaccharide and predictions of the composition of unknown capsule types (see below).

### Differentiation of K-types by FT-IR spectroscopy

FT-IR spectroscopy detects variation on the vibrational modes of chemical bonds that are exposed to infrared radiation, and when applied to bacterial cells provides a highly specific whole-organism fingerprint that reflects their biochemical composition. Hence, we evaluated the ability of this methodology to differentiate the 19 *Kp* capsule types or other surface structures (isolates with *wzi*151 or *wzi*200 were excluded since they included only 2 isolates each). We compared spectra from all corresponding isolates obtained in the same experimental conditions, and analyzed spectral variance by multivariate data analysis (see the section Methods for further details).

#### General features of FT-IR spectral data

FT-IR spectra of all *Kp* isolates displayed typical bacterial bands that had been previously related with the presence of different biomolecules such as lipids (W_1_, 3000–2800 cm^-1^), proteins/amides I and II (W_2_, 1700–1500 cm^-1^), phospholipids/DNA/RNA (W_3_, 1500–1200 cm^-1^), polysaccharides (W_4_, 1200–900 cm^-1^) and a fingerprint region (W_5_, 900–700 cm^-1^) ^31^. The highest spectral variance was detected in the region dominated by vibrations of carbohydrates (W_4_, 1200–900 cm^-1^) (data not shown). This region was selected for spectral data analysis. Considering that the capsule is the most variable surface structure and that it is mainly composed of polysaccharides, spectral diversity was analyzed and represented in supervised models considering K-types as classes. Two consecutive partial least squares discriminant analysis (PLSDA) models were used to obtain the highest level of correct predictions for all classes (15 in Model 1 and 7 in Model 2) (**Figure 2a, Figure 3a**). In these models we observed several well-defined clusters of isolates that were absolutely consistent with the K-type. To further evidence that FT-IR spectroscopy differentiation is based on K-type variation we developed an additional PLSDA model, now using STs as classes, instead of K-type (**Figure S1**). In this model, we observed also several well-established clusters that were superimposable with those obtained for K-types when one ST harbored only one K-type distinct from all the others identified (**Figure S1**).

**Figure 2.**
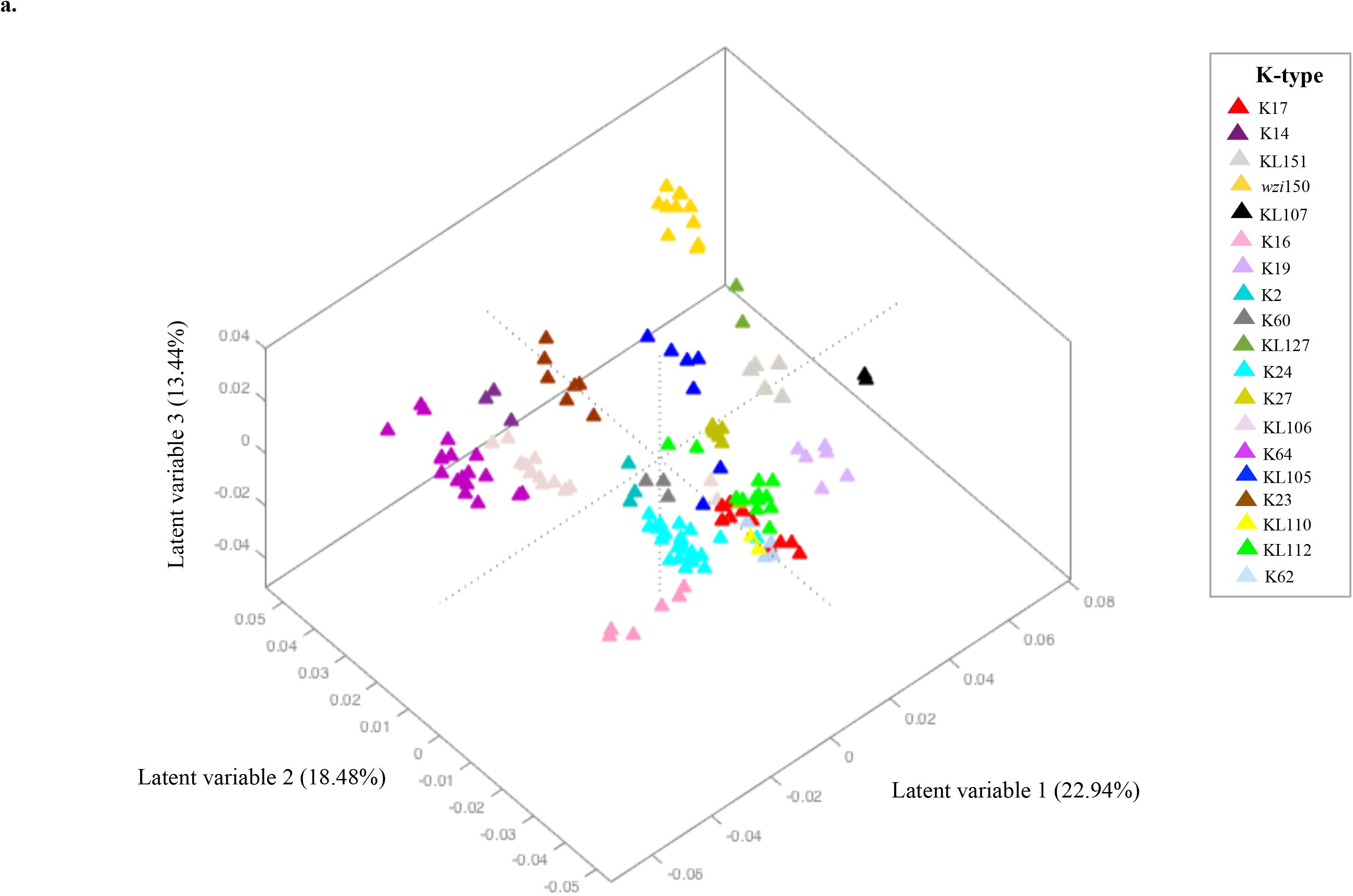
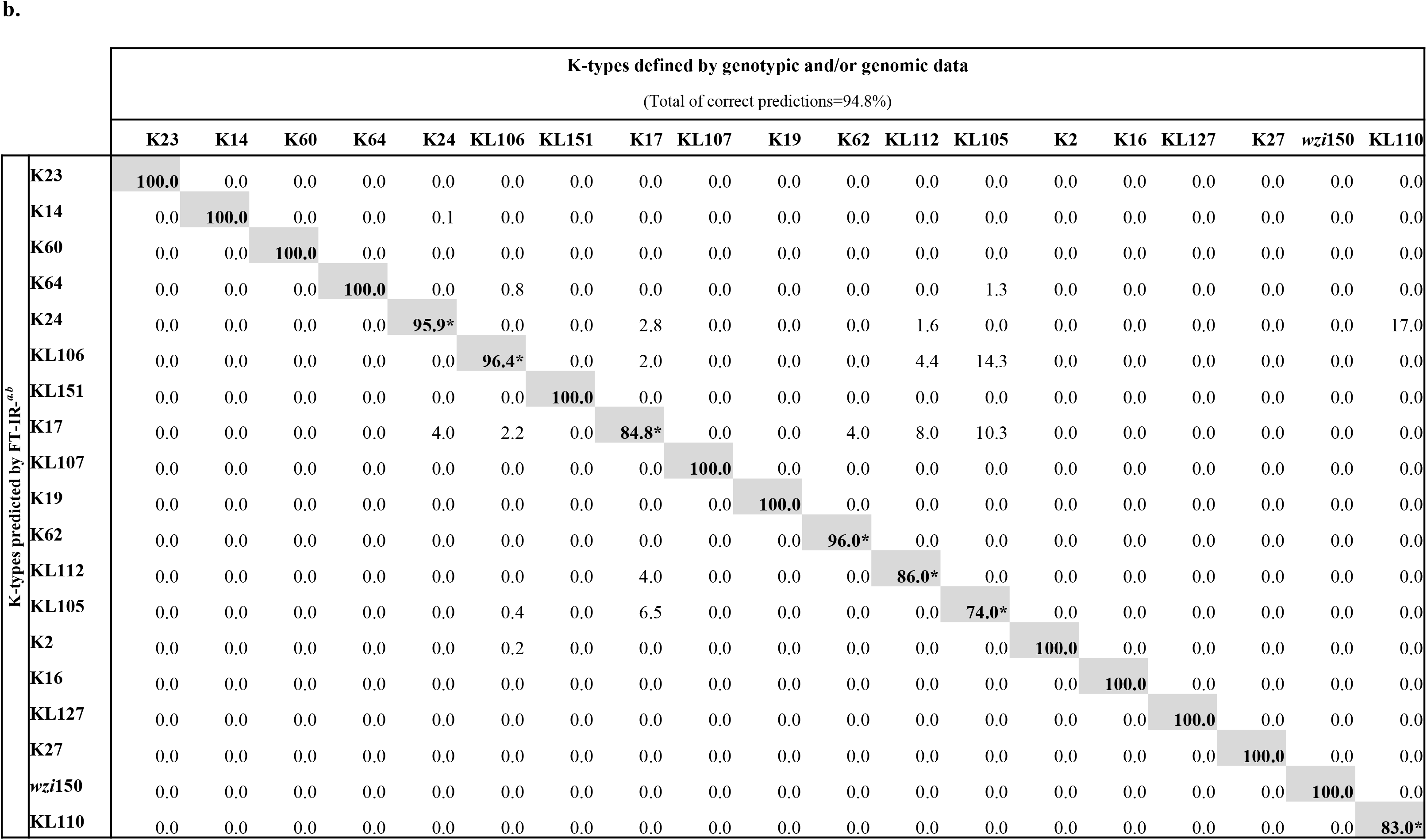
a Score plot of the PLSDA regression model 1 according to K-types corresponding to the first three latent variables (LVs). **Figure 2b.** Confusion matrix for *K. pneumoniae* PLSDA model 1 according to K-types defined by genotypic and/or genomic data (values are in %). Legend: ^*a*^ Grey shaded represent the percentage of isolates correctly predicted using FTIR-ATR for each K-type; ^*b*^ Values obtained considering 20 LVs in PLSDA model; * K-types predicted by FT-IR with less than 100% of confidence and resolved in Model 2.

**Figure 3.**
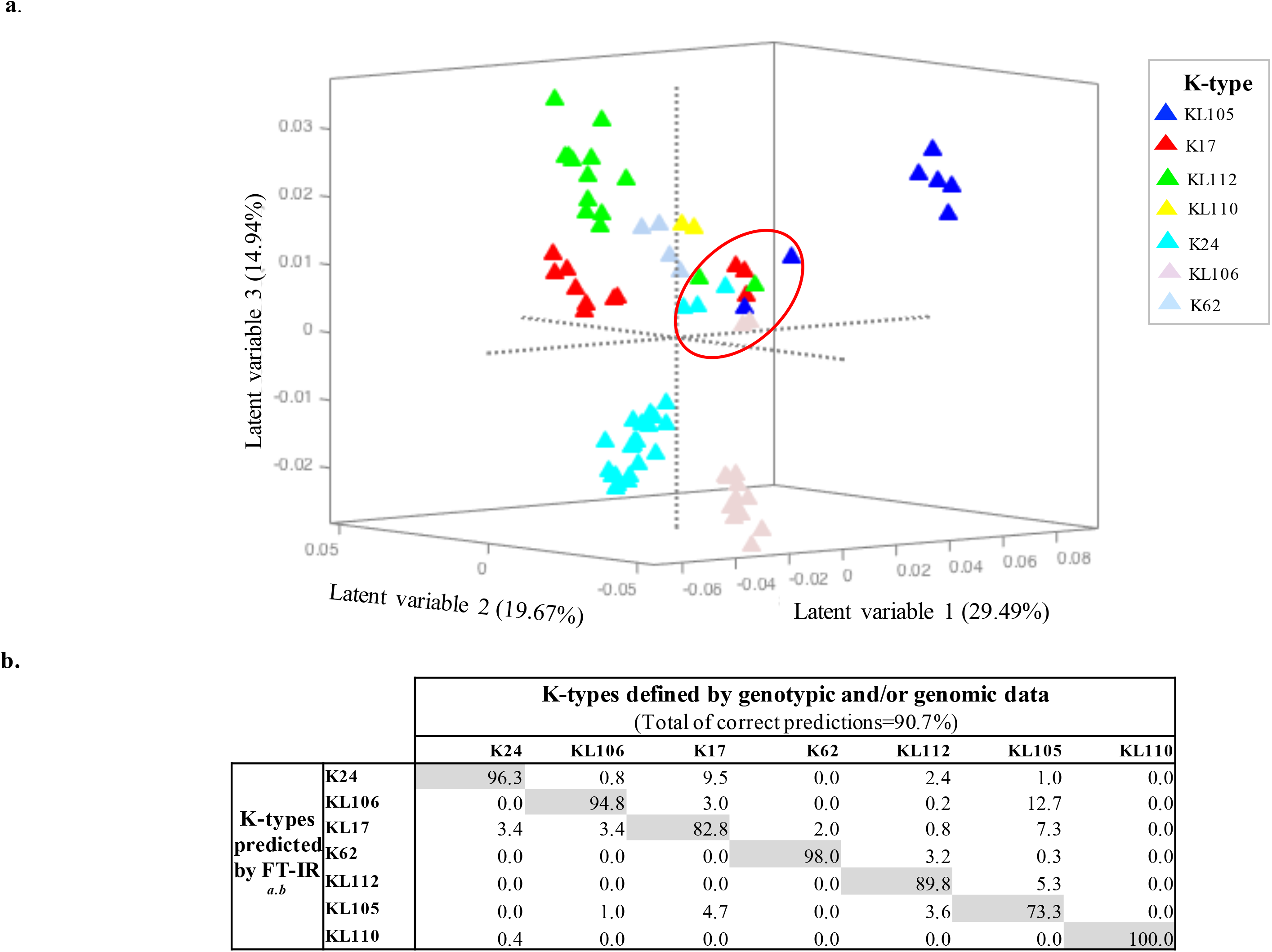
Score plot of the PLSDA regression model 2 according to K-types corresponding to the first three latent variables (LVs). Legend: The red circle includes the isolates that have a different phenotypic behavior from its main class**. Figure 3b.** Confusion matrix for *K. pneumoniae* PLSDA model 2 according to K-types defined by genotypic and/or genomic data (values are in %). Legend: ^*a*^ Grey shaded represent the percentage of isolates correctly predicted using FTIR-ATR for each K-type; ^*b*^ Values obtained considering 12LVs in PLSDA model;

#### Full K-type resolution in two PLSDA models

The analysis of PLSDA models considering K-types as classes (a different K-type was assumed for isolates exhibiting *wzi*150) (Model 1), fifteen clusters of isolates exhibiting 15 different K-types were perfectly distinguished with 94.8% of total correct K-types predictions (**Figure 2b**). These clusters included isolates with the same K-type from different STs (e.g. isolates from ST11/ST147 with K64) and also isolates belonging to O1 or O2 indistinctly (e.g. K64 isolates). In fact, O1 and O2 have highly similar structures that are most probably indistinguishable by FT-IR spectroscopy. They are both composed of galactose homopolymers (alternating β-D-Gal*f* and α-D-Gal*p* residues) named D-galactan-I (gal-I) or D-galactan III (gal-III) (O2 serogroup), that when capped with the gal-II (an antigenically different α-D-Gal*p* and β-D-Gal*p* disaccharide) forms the O1 serogroup. Classes that presented some cross-predictions and whose prediction rates were below 100% (n=7; K24, K17, K62, KL105, KL106, KL110, KL112), were modelled independently using a second PLSDA (Model 2). Results for Model 2 (**Figure 3a**) show that each seven K-types were distinguished and the proportion of correct predictions improved (0.4-17%) for several of them (**Figure 3b**). In fact, lower prediction rates were observed in heterogeneous classes that included *a priori* a few isolates that revealed a different phenotypic behavior (called “exception isolates”) than that expected for the respective class (**Figure 3a**; see below). A purge of isolates revealing differences at both genotypic and phenotypic level increased the correct prediction rates towards 100% (data not shown).

The FT-IR-based typing method discriminated the 19 different K-types tested, supporting differences in their final capsule composition or structure, including the biochemically uncharacterized KL-types. It provided a resolution identical to that of whole *cps* sequencing for discriminating closely related K-types (K14 and K64) or discrepant K-/KL-types (KL60, KL112). Also, our data suggests that *wzi*150 isolates are KL163 or KL46 since they were perfectly distinguished from KL27 isolates. Moreover, not only precise phenotypic-genotypic correlations were established, but also this methodology depicted phenotypic differences in a few isolates (8.9%; n=12/152, “exception isolates”) that were not predicted by genome data. Some were indistinguishable by *wzi*-based typing while others presented small differences in other sites of the locus that could be neglected (**Table 1**). For one third of them (n=4/12), it was possible to support the different phenotypic features by sequencing of *cps* regions other than *wzi*, as explained below.

#### Exploring capsular phenotypic – genotypic discrepancies

First, one KL112 isolate (ST17) incorrectly predicted might represent one of the few cases where differences in the O-type might impact on FT-IR spectra. This isolate was classified as O5, which, instead of D-galactan from O1 or O2, is composed by a homopolymer of mannose yielding a different polysaccharides’ configuration ^17,32^. Secondly, 2 KL105 ST11 isolates were distinguished by FT-IR spectroscopy from the others predicted as KL105 (**Figure 3a**). Main differences in the spectra from the two subgroups (arbitrarily designated as KL105-1 and KL105-2) were observed in the 1080-980 cm^-1^ region (spectral region dominated by ring vibrations of the carbohydrate region), suggesting variations in final capsule composition ^31^ (**Figure 4a**). They all share the same PFGE-type and O-type (O2) and were isolated in Portugal as DHA-1 or DHA-6 producers during a large period of time (2006-2013) (**Table 1**) (Gonçalves et al., unpublished results). The whole *cps* operon (27 kb in size) of 2 representative isolates (1 KL105-1 and 1 KL105-2) was extracted from whole genome sequences and compared. A 10 bp deletion was found in the *wzi* of KL105-2 isolate that is outside from the region sequenced by *wzi*-based typing, resulting in a premature stop codon and a Wzi protein with 427aa instead of 477aa (**Figure 4b**). Considering that the Wzi protein is involved in the assembly of the capsule, it is possible that changes in this assembly might affect the final capsule composition or amount ^33^. Third, one K24 isolate (H1119) predicted by *wzi* sequencing had recombinant K24/K39 *cps* locus (sequence was deposited in GenBank database with accession number NXBL00000000). Fourth, FT-IR detected differences in 2 KL106 isolates. An *in silico* analysis of 496 ST258 genomes by Kaptive (publicly available at NCBI) revealed that 153 of them (31%) carried *wzi*29 that differed in the presence or absence of IS sequences (IS1-IS3, +1588bp) in specific sites of the *cps* locus, suggesting the circulation of two KL106 variants. Thus, we hypothesize that the two clusters depicted by FT-IR for KL106 might represent these two variants (data not shown).

**Figure 4.**
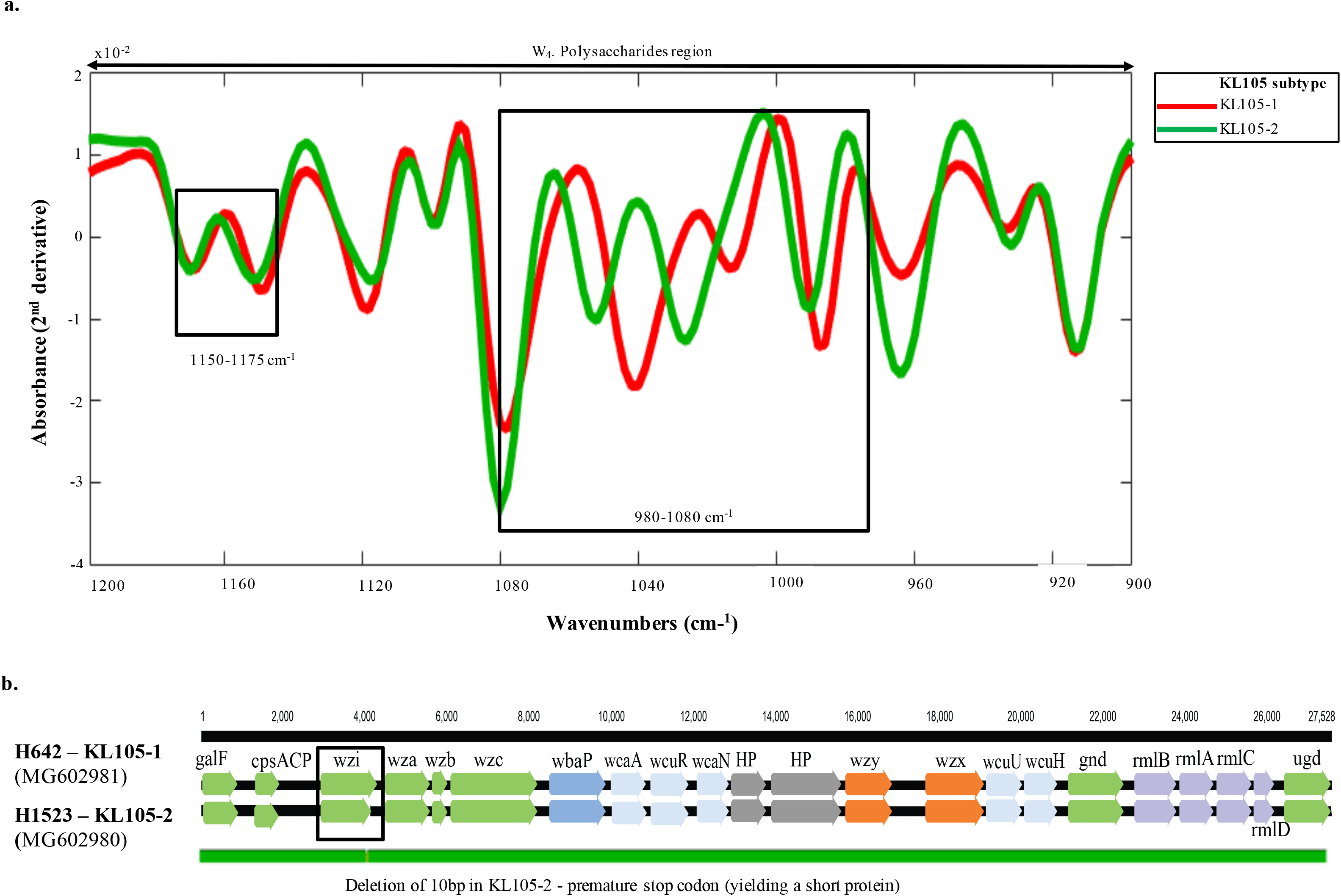
a *K. pneumoniae wzi*75-KL105 FT-IR spectra processed with SNV and Savitzky-Golay (9 points filter size. 2^nd^ degree polynomial. 2^nd^ derivative) corresponding to the mean spectra in the region 900-1200 cm^-1^. **Figure 4b**. Representation of *cps* genetic cluster of KL105-1 and KL105-2 isolates.

Thus, our data demonstrates differences in capsule composition of main *Kp* K-types can be reliably detected by FT-IR spectroscopy, which provides a resolution identical or even greater than that of the most discriminatory genotypic-based K-typing method (WGS).

### Correlation between FT-IR K-types and capsule biochemical composition

To unequivocally settle the basis for FT-IR-based K-type discrimination, we supported the phenotypic differences detected by FT-IR spectroscopy with the chemical composition available for capsule types. For this purpose, we represented the similarity of the spectra in a dendrogram generated by hierarchical cluster analysis (HCA) and correlated the assignments with the chemical composition of the different known K-types (**Figure 5a**). Using this unsupervised method, the nineteen K-types modeled in Figures 2a and 3a were also discriminated in clusters defined at distances <0.4, that grouped isolates according to their K-type in accordance with clusters defined in PLSDA models. In parallel, we represented in **Figure 5b** the composition and structure of 12 out of 19 known K-types (including the newly described KL107), using polysaccharide composition and structural data obtained previously for one representative strain from each K-type (**Table S1**). We observed that these capsular types exhibit a marked diversity of patterns based on the size of the polysaccharide polymer, the number and type of monosaccharides, the type of linkages or the presence of side chains or modifications of the lateral sugars, that are on the basis of correct FT-IR-based discrimination. They vary between tetra- and heptasaccharides made-up of glucose, glucuronic acid, mannose, rhamnose, fucose, galactofuranose or galacturonic acid in different proportions and order, though some appear to have similar structures (**Figure 5b**).

**Figure 5.**
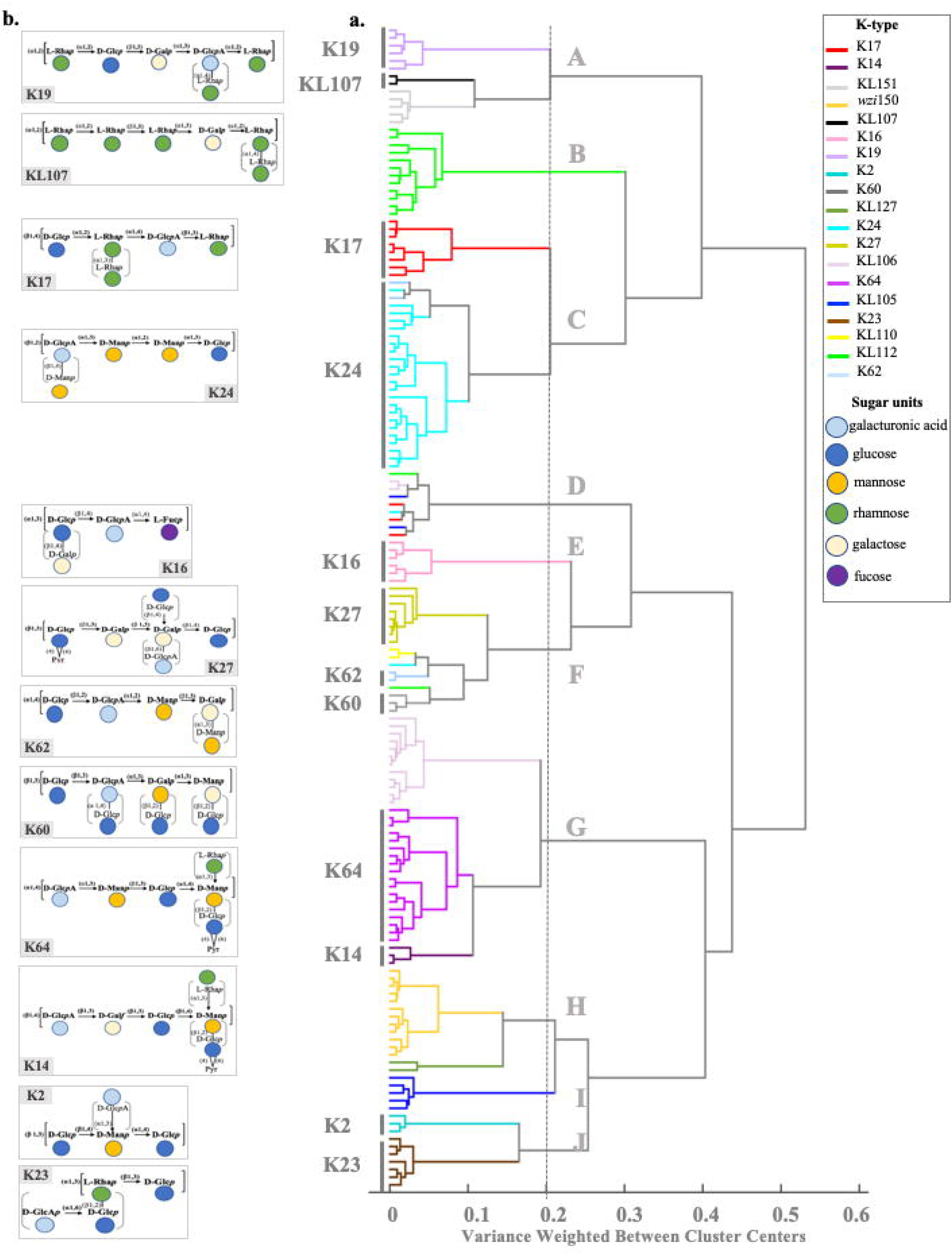
a Dendrogram obtained from the 1200–900 cm^-1^ spectral region using the Ward’s algorithm and 13 principal components (PCs) distance for isolates of the different K-types. **Figure 5b.** Known CPS chemical structures of *K. pneumoniae* included in this study. Legend: D-Gal, D-galactose; D-Galf, galactofuranose, D-Glc, D-glucose; D-Man, D-mannose; L-Rha, L-rhamnose; L-Fuc, L-Fucose, Pyr, pyruvate; Square brackets represent a single K-unit structure. Arrows represent the type of linkages between the monosaccharides within the K-unit. Curved brackets represent additional monosaccharides and/or pyruvate groups in the K-unit structures.

#### Analysis of similar K-types inferred from FT-IR spectra

A high similarity between K-types K19 and KL107 (**Figure 5a** branch A), K17 and K24 (**Figure 5a** branch C), K14 and K64 (**Figure 5a** branch G), and K2 and K23 (**Figure 5a** branch J) is inferred from the HCA (distances below 0.2), which is supported by their closely related K-type structures, as explained below (**Figure 5b).**

*K19 and KL107*. These capsular types are both composed by a hexasaccharide that have in common a high number of rhamnose residues (3 and 5, respectively), and vary slightly in the composition of other sugars. Whereas K19 contains a polymer of D-galactose, D-glucose, L-ramnose (3 monomers) and D-glucoronic acid, KL107 contains D-galactose, D-galacturonic acid and rhamnose (4 monomers).

*K17 and K24*. These capsular types consist on a similar pentasaccharide structure, containing D-glucose, D-glucuronic acid and 3 L-rhamnose (K17) or 3 D-mannose (K24), that differ only in the final hydroxyl group.

*K14 and K64*. They are composed of highly similar hexasaccharides composed by D-glucose (2 monomers, one of them acetylated), D-glucuronic acid, L-rhamnose and either 2 D-mannoses (K64) or 1 L-mannose and 1 galactofuranose (K14). In fact, both structures are highly similar even in configuration and these K-types yield cross-reactions in serological methods.

*K2 and K23*. These two capsular types are characterized by tetrasaccharides in different configurations, composed of D-glucose (2 monomers) and D-glucuronic acid and either D-mannose (K2) or L-rhamnose (K23) that are highly similar sugars differing only in the conformation and the final group (CH_3_ or CH_2_OH).

Additionally, KL60, KL62, K27 were all grouped in branches F from Figure 5a, that included also a few K-types for which the structure is not known, and for this reason any comparison lacks robustness. We observed that KL60, KL62, K27 are diverse in structure (penta-heptasaccharides) and composition (variable but especially enriched in glucose). K16 appears in a separate branch (E) from the dendrogram and is clearly distinguished from all the others, since it is formed by a tetrasaccharide containing D-glucose, D-glucuronic acid, D-galactose and it is the only one containing L-fucose.

The correlations established strengthen FT-IR-based K-type assignments and highlight the need to both characterize the structure/composition of new KL-types and increase reliability of the clustering and the comparisons with a higher number of isolates from certain K-types.

### Prediction of the capsular composition based on FT-IR spectroscopy assignments

Several K-types included in this study are linked to worldwide spread *Kp* lineages (KL105, KL106, KL110, KL112, KL127, KL151) encountered among MDR *K. pneumoniae* clinical isolates for which the structure has not yet been characterized. In this section, we provide insights into the possible structure and composition (type of sugars) for these new KL-types considering the similarity between FT-IR spectra (**Figure 5a**) combined with the information contained in their corresponding *cps* locus (**Figure 1**).

Since spectra obtained from isolates exhibiting KL105 and KL127 clustered with K2 and K23 types (distance<0.3), we predict a tetrasaccharide structure composed of D-glucose, D-glucuronic acid and possibly D-galactose and D-rhamnose, that is also supported by the presence of genes *wbaP* and *rmlBADC*, respectively, in the *cps* operon. KL112 capsule is closely related with that of K17 or K24 at a distance >0.3. Thus, we predict that it could be composed of a pentasaccharide of D-glucose, D-galactose and D-mannose, which is also corroborated by the presence of *wbaP* and *manCB* on the *cps* cluster. Similarly, KL151 capsular type is highly related with KL107 and thus we expect to be a hexapolysaccharide composed of several rhamnose residues (supported by the presence of *rmlBADC*) and absence of D-galactose (*wcaJ* instead of *wbaP*). KL110 might be a pentasaccharide comprising several units of D-glucose and/or D-galactose and mannose according to the corresponding operons encountered in the *cps* operon.

Thus, using our FT-IR-based framework, we predicted for the first time the presumptive structure/composition of new KL-types, that was supported on *cps* genotypic data. Further studies are needed to validate these predictions and potentiate the use of FT-IR spectroscopy as a phenotypic method for K-type identification and characterization.

## DISCUSSION

In this study, we establish for the first time a framework to support FT-IR spectroscopy as a quick, reliable and inexpensive phenotypic method for identification of *Kp* capsular types. The multidisciplinary strategy used allowed clarifying the fundamentals for FT-IR-based K-type discrimination, increasing knowledge on K-type phenotypic variation (especially on the newly described KL-types), and validating the methodology for K-typing, outbreak management and epidemiological surveillance of MDR *Kp*.

We demonstrated that FT-IR-based K-type differentiation relies on the distinctive biochemical profiles obtained in the spectral region dominated by carbohydrates (W_4_, 1200-900 cm^-1^) ^31^. It was further substantiated with whole *cps* locus analysis and correlated with the number and type of monomers that make up the capsular polysaccharide, whereas the order and type of bonds (alpha or beta) does not seem to influence K-type prediction. The same spectral region was previously reported to be highly discriminatory for several bacterial species including other relevant clinical or food pathogens such as *Escherichia coli, Acinetobacter baumannii, Salmonella enterica, Staphylococcus aureus* or *Streptococcus pneumoniae* ^24–26,34,35^. Some of these studies have also pointed-out a correlation with variation on bacterial serogroups or capsules, supporting the importance of surface polysaccharides as discriminatory biomarkers ^22^.

In fact, the importance of surface structures (and especially the capsule on capsulated bacteria) on evolution, pathogenesis and host adaptation of bacterial pathogens is well known. However, full understanding on K-type variation has been hindered by variable drawbacks of available methods for K-typing, and reawakened only with the burst of genotypic or genomics-based approaches ^17,36^. Nevertheless, the putative number of capsule types encountered varies strongly with the method used. Up to now, there have been recognized 557 *wzi* and 120 *wzc* alleles (http://bigsdb.pasteur.fr/klebsiella/klebsiella.html), the latter probably less affected by recombination events. On the other hand, 161 *cps* loci (KL-) types have been predicted by whole *cps* sequencing, which is twice the number of K-types initially recognized by traditional serotyping techniques (77 K-types) ^7^. In the absence of structural data on these new K-types, it remains to be clarified if all of them correspond to biochemically distinct types and their correlation with the 77 K-types of reference. Our FT-IR-based approach supports a specific capsular structure for each one of the K-types tested, including many of the new K-types identified among clinically relevant MDR clones, and also closely related ones (e.g. K14 and K64). In this sense, FT-IR spectroscopy could be extremely useful in the phenotypic validation of the new inferred KL-types and guide the selection of those to be characterized biochemically, a highly desired goal ^36^. In addition, our FT-IR-based approach unveiled phenotypic differences within isolates considered from the same class (e.g. KL105 or KL106), whose significance is unclear when only genomics data is considered, reinforcing the sensitivity of the method. In fact, FT-IR-based assignments suggested that these “exception isolates” should indeed be regarded as independent classes for which a different capsular biochemical composition is predicted. The frequency with which these phenomena occur in the clinical setting, the reasons underlying or even their significance in the context of host interactions are unknown, but they seem to occur at least occasionally ^37^.

Currently, the correct prediction of K-types in *Kp* depends on a combined approach of genetic markers, epidemiological data and/or comparative genomics of *cps* loci, which is not straightforward, requires expertise and there is a gap between genetically- or serologically-defined K-types. Our data strongly supports FT-IR-based K-typing as a highly reliable alternative method that might represent a promising K-typing diagnostic tool. In addition, the ability of FT-IR spectroscopy to discriminate *Kp* capsular types represents a major advantage since K-types were found to be good epidemiological markers of particular *Kp* lineages with biological significance. The link between a particular clone and the K-type had been already recognized for HV *Kp*, but much less was known regarding the widespread MDR *Kp* population ^5^. Detailed and comprehensive phylogenomics studies have shown a high specificity between certain lineages and particular K-types within CG11, CG14, CG15 or CG258 ^7,14,15,19,20,28^. In fact, MDR *Kp* expansion causing hospital or community acquired infections has been driven by clonal selection, since the population is dominated by a small number of lineages exhibiting particular K-types, where phenomes of capsular switching seem to occur sporadically at least in short time-scales (this study, Kaptive database) ^6,7,38,39^. In some cases, capsular recombination events involving phenotypic changes might represent important evolutionary steps with several biological consequences, as occurred with the two different ST258 clades exhibiting KL106 or KL107 K-types, that are of interest to detect ^40,41^. It was based on this information that we selected representative widespread MDR lineages characterized at genomics and/or molecular level to include in this study, and showed that they were correctly depicted by FT-IR spectroscopy, reflecting the high discriminatory power of the methodology.

Besides K-types differentiation, our in-house FT-IR *Kp* models are being used routinely and successfully to classify unknown *Kp* isolates, where over 500 isolates from different hospitals, long-term care facilities, community laboratories have already been tested (data not shown). The spectra obtained from new isolates are compared with our own databases and their projection on the PLSDA models generated in this study provides a tentative K-type assignment, that is being corroborated by *wzi* sequencing. On this basis, the method has been crucial in the recognition and early detection of several hospital outbreaks involving carbapenemase (KPC-3, OXA-48, NDM-1), ESBL and/or MCR-1 producers, where in some cases useful information (K-type, tentative clone assignment according to local epidemiology) is provided in 24h ^42,43^ (Novais et al. unpublished data). This methodology is simple and inexpensive since only one fresh colony is needed (see Material and Methods for further detail), the cost of the equipment is significantly lower than that of other competitors (6 times less than MALDI-TOF MS or 3 times less than most common sequencing platforms) and the cost of its maintenance is very low. Furthermore, reproducible results were obtained at least in the same instrument and in FT-IR equipment’s from different manufacturers (Frontier from Perkin-Elmer and FT-IR Alpha from Bruker), and also using variable experimental conditions (+/-4h incubation time, different culture media) (data not shown), assuring the stability of the method in variable environmental conditions. We recognize that its usefulness for routine microbiology laboratories depends on the adaptation of the method for a non-specialist user, that depends on the creation of judicious databases created under standardized conditions and automation of data analysis. Indeed, the potential of the methodology has already been recognized by Bruker, that launched a FT-IR-based equipment (IR Biotyper®) in June 2016, intended to be used for routine strain typing using a simple and automated process.

## CONCLUSIONS

In this study, we demonstrate an unprecedent resolution at a fast and low-cost rate of *Kp* K-types at the phenotypic level, supporting FT-IR spectroscopy as a reliable cost-effective tool for routine K-typing and for validation of genomics-based K-type predictions. By providing a resolution identical or even higher than whole genome sequencing, it can be an asset for outbreak management and also surveillance in well-known epidemiological contexts. Finally, the multidisciplinary strategy used supports that our FT-IR-based approach might be extremely useful to improve our understanding on sugar-based coating structures of high relevance for strain evolution and host adaptation of a key bacterial pathogen.

## MATERIAL AND METHODS

### Bacterial Strains

One-hundred fifty-four well-characterized MDR *Kp* clinical isolates representing main Clonal Groups (CG) circulating in different geographic regions (Brazil, Greece, Poland, Portugal, Romania, Spain) for long periods of time (2002-2015) were selected to validate the approach. Clonal relatedness among the isolates was evaluated by gold-standard and reference genotypic methods (multilocus sequence typing and pulsed-field gel electrophoresis), depicting 13 STs (representing 7 CG) and 35 PFGE-types. Most of the isolates were producers of ESBLs, acquired AmpCs and/or carbapenemases, and were enriched in particular virulence factors, such as the urease cluster (100%), type 1 and 3 fimbriae (99.4%), the yersiniabactin siderophore (*ybtS*) and iron transporter permease genes (*kfuBC*) (64% and 37%, respectively), the latter with variable distribution in the collection analysed. Details about the bacterial isolates included in this study are summarized in **Table 1**.

### Genotypic and phenotypic characterization of surface polysaccharides structures

In all isolates, PCR and sequencing of specific genetic markers were used for genotyping of K- and O-types. For the genotypic-based prediction of K-types, we sequenced a 447 bp fragment from a highly variable region of the *wzi,* and occasionally specific *wzy* fragments ^14,15^. Regarding O-genotyping, specific regions of *wzm* and *wzt* genes from the *rfb* cluster were amplified for O1/O2, O3 and O5 identification. Furthermore, and additional PCR as performed to distinguish O1 and O2 and its variants (designed in the *wbbY* loci unlinked to the *rfb* cluster) ^10,17^. Additionally, WGS was performed for 8 isolates for which discrepancies between genotypic and phenotypic features were observed. WGS was performed by Illumina MiSeq (2x300bp pair-ended runs, ∼6Gb per genome, coverage 100x), and reads assembled using SPAdes version 3.9.0 (http://bioinf.spbau.ru/en/spades) and full *cps* locus further annotated with Geneious R10 software considering the nomenclature proposed by Reeves et al. ^44^. The sequences of the complete *cps* operon were deposited at GenBank database under the accession numbers MG602975 to MG602982 and under the BioProject PRJNA408270.

Phenotypic characterization of surface bacterial components was performed using FT-IR spectroscopy with attenuated total reflectance (ATR) mode, as previously described ^34,35^. Briefly, isolates were grown on Mueller-Hinton agar at 37°C for 18h and colonies were directly transferred from the agar plates to the ATR crystal and air-dried in a thin film. Spectra were acquired using a Perkin Elmer Spectrum BX FT-IR System spectrophotometer in the ATR mode with a PIKE Technologies Gladi ATR accessory from 4000–600 cm^-1^ and a resolution of 4 cm^-1^ and 32 scan co-additions. For each isolate, at least three instrumental replicates (obtained from the same agar plate in the same day) and three biological replicates (obtained in three independent days) were acquired and analysed, corresponding to a minimum of nine spectra per strain ^35,45^.

### Spectral data analysis

All chemometric analyses was performed using Matlab R2015a version 8.5 (MathWorks, Natick, MA) and PLS Toolbox version 8.5 for Matlab (Eigenvector Research, Manson, WA, USA). Original FT-IR spectra were processed with standard normal variate (SNV) followed by the application of a Savitzky-Golay filter (9 smoothing points, 2^nd^ order polynomial and 2^nd^ derivative) ^46,47^. Prior to modelling with PLSDA, spectra were mean-centred. Due to the high amount of generated data, a mean spectrum of each isolate (resulting from at least nine congruent replicates) was considered in the analysis performed. Spectra were analyzed by a supervised (partial least squares discriminant analysis, PLSDA) chemometric model using for discriminatory purposes the region of the spectra corresponding to the carbohydrates vibrations (W4, 1200-900 cm^-1^) ^31^. PLSDA is a supervised method based on the PLS regression method. In PLSDA models, we assign to each isolate spectrum (*x*_*i*_) a vector of zeros with the value one at the position corresponding to its class (*y*_*i,*_ ST or K-type), in such a way that categorical variable values (*y*_*i*_) can be predicted for samples of unknown origin. Model loadings and the corresponding scores were obtained by sequentially extracting the components or latent variables (LVs) from matrices *X* (spectrum) and *Y* (matrix codifying K-types). In PLSDA, a probability value for each assignment is estimated for each sample. The number of latent variables (LVs) was optimized using the leave-one-sample-out cross-validation procedure in order to prevent over-fitting considering only 70% of the available data (randomly selected). After optimization of the number of LVs, the model was tested on the remaining 30% samples in order to assess the proportion (%) of correct predictions for each class ^35,45,48^.

The unsupervised method hierarchical cluster analysis (HCA) was also applied to evaluate the spectral similarity between isolates (and eventually to correlate clusters with K-type structures). The dendrograms were obtained using the Ward’s algorithm and a preliminary principal component analysis to ensure the robustness of the results. Thirteen components were retained. The same preprocessing and scaling used for PLSDA was used for HCA.

## ACKNOWLEDGEMENTS

We thank Marek Gniadkowski (National Medicines Institute, Poland), Vivi Miriagou (Department of Bacteriology, Hellenic Pasteur Institute, Athens, Greece), Grigore Mihăescu (University of Bucharest, Faculty of Biology, Deptartment of Microbiology, Bucharest, Romania), Rafael Cantón and Teresa Maria Coque (Laboratorio de Microbiologia, Hospital Universitario Ramón y Cajal, Madrid, Spain), Leonardo Neves de Andrade and Ana Lucia Darini (Universidade de São Paulo, Faculdade de Ciências Farmacêuticas de Ribeirão Preto) for the strains analyzed in this study.

## Supplementary Material

**Figure S1.** Score plot of the PLSDA regression according to STs. corresponding to the first two latent variables (LVs).

**Table S1.** Reference capsular (K) types included in this study

