## Supplementary material for "On the front line of *Klebsiella pneumoniae* surface structures understanding: establishment of Fourier Transform Infrared (FT-IR) spectroscopy as a capsule typing method": Figure S1

**Figure S1**. Score plot of the PLSDA regression according to STs. corresponding to the first two latent variables (LVs).

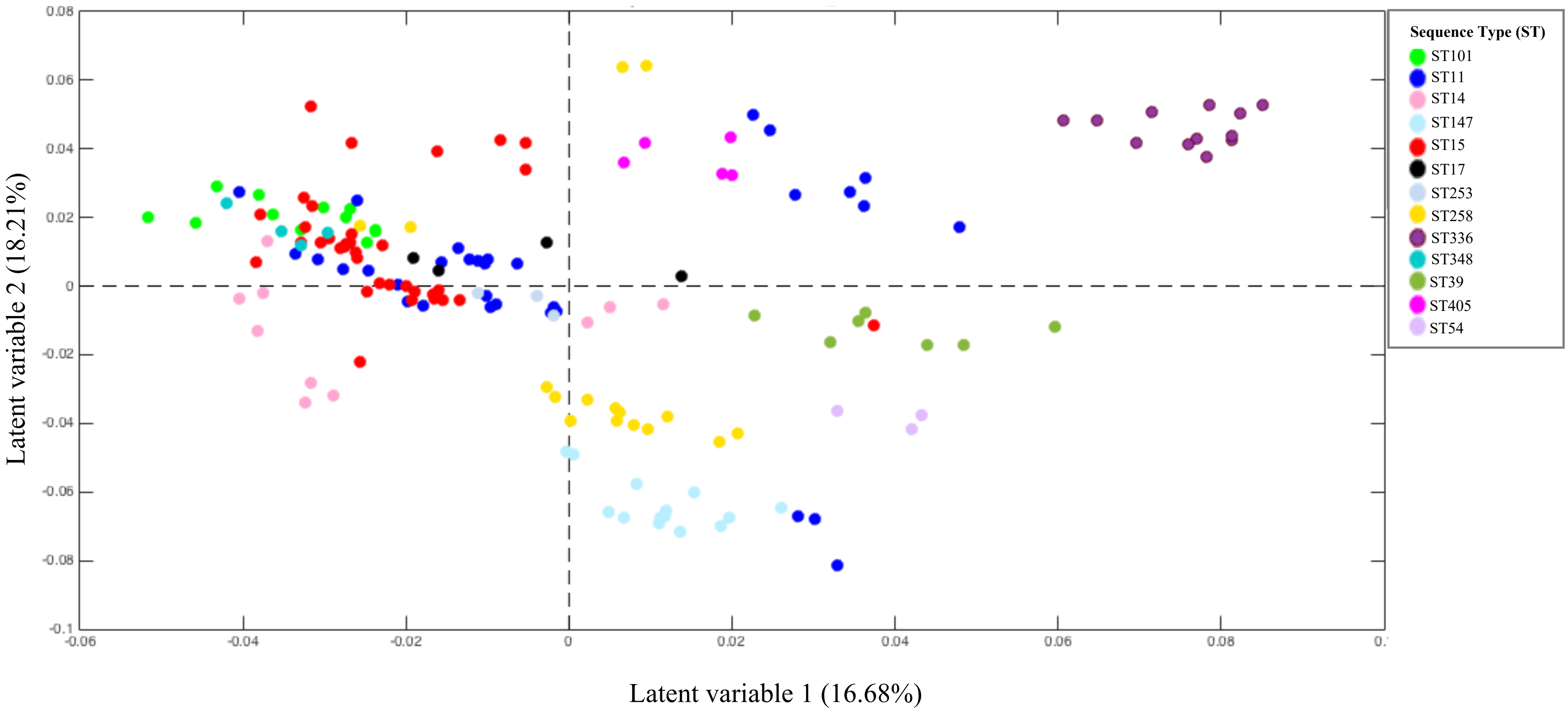
