## Supplementary material for "On the front line of *Klebsiella pneumoniae* surface structures understanding: establishment of Fourier Transform Infrared (FT-IR) spectroscopy as a capsule typing method": Table S1

**Table S1**. Reference capsular(K) types included in this study

| **Capsular type** | **Strain** | **GenBank Accession nr.** | **Reference** |
| --- | --- | --- | --- |
| **K2** | VGH525 | AB371296 | ^1^ |
| **K14** | VGH916 | AB371294 | ^2^ |
| **K16** | 2069/49 | AB742228 | ^3^ |
| **K17** | 2005/49 | AB924557 | ^4^ |
| **K19** | 293/50 | AB924559 | ^5^ |
| **K23** | 2812/50 | AB924561 | ^6^ |
| **K24** | 1680/49 | AB924562 | ^7^ |
| **K27** | 6613 | AB924565 | ^8^ |
| **K60** | 4463/52 | AB924597 | ^9^ |
| **K62** | VGH698 | AB371295 | ^10^ |
| **K64** | NCTC 8172 | AB924600 | ^11^ |
| **KL107** | 2796 | GCA_000281375.1 | ^12^ |
